# Comparative Analysis of Feature Selection Methods for Single-Cell RNA Sequencing Data

**DOI:** 10.64898/2025.12.02.691907

**Authors:** Adham M. Alkhadrawi, Mohammed A.B. Mahmoud, Mian M.Y. Khalil, Abdullah All Jaber

## Abstract

Feature selection is a critical preprocessing step in single-cell RNA sequencing (scRNA-seq) analysis, directly impacting downstream clustering and biological interpretation. We systematically compared 16 feature selection methods across three diverse datasets: PBMC3K (immune cells), Visium Heart, and Visium Brain (spatial transcriptomics). Methods included established approaches (Seurat HVG variants, CellRanger HVG), statistical methods (Pearson residuals, variance-based, coefficient of variation), supervised methods (ANOVA F-test, mutual information, random forest), and deep learning techniques (IntegratedGradients, DeepLIFT, GradientShap). We evaluated methods based on execution time, feature overlap, marker recovery, and pathway enrichment. Our analysis revealed substantial variability across methods, with mean pairwise overlap of only 23.7%. However, a core set of 1,150 genes was consistently selected by *≥* 50% of methods. Supervised methods demonstrated superior recovery of known cell type markers, while unsupervised approaches captured broader biological processes. Deep learning methods identified unique gene sets with strong immune pathway enrichment but higher computational cost. Pathway analysis showed all methods successfully identified relevant biological processes, though with varying emphasis. These findings provide practical guidance for method selection based on analytical goals and highlight the value of ensemble approaches in scRNA-seq feature selection.

## 1 introduction

Single-cell RNA sequencing (scRNA-seq) has revolutionized our understanding of cellular heterogeneity, enabling the exploration of gene expression profiles at an unprecedented resolution [1, 2]. This technology has become indispensable in fields ranging from developmental biology to cancer research, providing insights into cellular differentiation, immune responses, and disease mechanisms [3, 4]. However, the high dimensionality and sparsity of scRNA-seq data present significant analytical challenges, particularly in the critical step of feature selection [5, 6]. Feature selection, the process of identifying the most informative genes for downstream analysis, is essential for reducing noise, improving clustering accuracy, and enabling biological interpretation [7, 8].

The importance of feature selection in scRNA-seq analysis cannot be overstated. It directly impacts the identification of cell types, the reconstruction of developmental trajectories, and the discovery of biomarkers for disease [9, 10]. Traditional methods for feature selection, such as highly variable gene (HVG) detection, have been widely adopted due to their simplicity and efficiency [11, 12]. However, these methods often struggle to capture the complex biological variation inherent in single-cell data, leading to suboptimal clustering and misclassification of cell types [13, 14]. Moreover, the performance of these methods can vary significantly across different datasets, highlighting the need for more robust and adaptive approaches [15, 16].

Recent advances in deep learning interpretability have introduced neural network attribution methods for feature selection. These techniques—IntegratedGradients [17], DeepLIFT (Deep Learning Important FeaTures) [18], and GradientShap [19]—train classifiers to distinguish cell types, then use gradient-based attribution to identify genes that most influence predictions. IntegratedGradients integrates gradients along a path from baseline to input, DeepLIFT compares activations to reference states, and GradientShap combines gradients with Shapley values from game theory. While computationally more expensive than statistical methods, these approaches may capture complex non-linear gene interactions that variance-based methods miss. However, their practical utility for scRNA-seq feature selection relative to established methods remains unclear, motivating their inclusion in this systematic comparison.

Despite these advancements, several challenges remain in the field of feature selection for scRNA-seq data. One major limitation is the lack of consensus on the optimal set of features for different biological contexts [5, 13]. While some methods excel in specific applications, such as cell type identification or trajectory inference, their generalizability across diverse datasets is often limited [11, 14]. Furthermore, the integration of multi-omics data and the incorporation of prior biological knowledge into feature selection algorithms are still underdeveloped, offering opportunities for future innovation [20, 21].

Emerging trends in feature selection include the use of hierarchical and context-dependent strategies to improve the accuracy and interpretability of selected features [7, 9]. For example, hierarchical marker gene selection strategies group similar cell clusters and select marker genes in a hierarchical manner, providing a more nuanced understanding of cell type relationships [7]. Additionally, the integration of single-cell and bulk RNA sequencing data, as demonstrated in studies on osteoporosis and osteosarcoma, has revealed immune-related biomarkers that could inform personalized treatment strategies [22, 14].

The rationale for this study is to address the critical gaps in current feature selection methods for scRNA-seq data. By conducting a comprehensive benchmark of 13 feature selection methods across three diverse datasets, we aim to provide practical guidance for method selection and highlight the need for context-dependent strategies [1, 2]. Our analysis includes established methods such as HVG selection variants from Seurat [23], CellRanger [24], and Scanpy [25], and machine learning-based methods via Scikit Learn [26]. We evaluate both computational performance and biological relevance, revealing distinct clustering patterns and substantial dataset-dependent selection patterns. Pathway enrichment analysis further demonstrates systematic method-specific biological biases, emphasizing the importance of selecting appropriate feature selection strategies for specific biological questions.

In conclusion, the field of feature selection for scRNA-seq data is rapidly evolving, with new methods and approaches continuously being developed to address the challenges posed by high-dimensional and sparse datasets. This study contributes to this ongoing effort by providing a comprehensive benchmark of existing methods and highlighting the need for context-dependent strategies. Our findings have important implications for the design of future studies and the development of more effective tools for single-cell data analysis.

## 2 Methods

### 2.1 Datasets

We analyzed three publicly available datasets representing different tissue types and sequencing technologies. The PBMC3K dataset (Figures S1 and S2) contains 2,700 peripheral blood mononuclear cells from a healthy donor, sequenced using the 10x Genomics Chromium platform. This dataset includes diverse immune cell populations including T cells, B cells, NK cells, monocytes, and dendritic cells. After quality control filtering, we retained 2,700 cells for analysis.

The Visium Heart dataset (Figures S3 and S4) comprises spatial transcriptomics data from human heart tissue (V1_Human_Heart section), containing 4,236 spatial spots. Each spot captures RNA from multiple cells, providing spatial context for gene expression patterns across tissue architecture. The Visium Brain dataset (Figures S5 and S6) contains spatial gene expression from human brain cortical tissue (V1_Human_Brain_Section_1) with 4,906 spots, enabling analysis of brain region-specific marker detection.

All datasets were downloaded using the Scanpy library and underwent standardized preprocessing. For each dataset, we generated two data layers: raw count matrices for methods requiring unnormalized data, and log-normalized expression matrices (normalized to 10,000 counts per cell, followed by log1p transformation) for methods expecting processed data. Quality control metrics were calculated including total UMI counts per cell, number of detected genes, and mitochondrial gene percentage. Cells with fewer than 200 detected genes or more than 20% mitochondrial content were excluded from the PBMC3K dataset. For spatial transcriptomics datasets, all spots were retained to preserve tissue architecture.

### 2.2 Clustering Preprocessing

To ensure consistency across all feature selection methods and enable fair comparison of supervised approaches, we performed Leiden clustering on each dataset before applying feature selection methods. For PBMC3K, we identified 11 clusters with sizes ranging from 11 to 532 cells per cluster (cluster distribution: 0: 532, 1: 437, 2: 356, 3: 344, 4: 343, 5: 173, 6: 154, 7: 150, 8: 149, 9: 51, 10: 11 cells). For Visium Heart, we created 9 clusters ranging from 235 to 674 spots per cluster (cluster distribution: 0: 674, 1: 639, 2: 621, 3: 585, 4: 529, 5: 376, 6: 331, 7: 246, 8: 235 spots). For Visium Brain, we obtained 11 clusters ranging from 308 to 641 spots per cluster (cluster distribution: 0: 641, 1: 574, 2: 561, 3: 540, 4: 463, 5: 439, 6: 380, 7: 355, 8: 336, 9: 309, 10: 308 spots). These cluster assignments served as cell type labels for supervised feature selection methods.

### 2.3 Feature Selection Methods

We compared 16 feature selection methods representing four broad categories:

#### Statistical Methods (9 methods)

- Seurat HVG: Models mean-variance relationship using local polynomial regression (scanpy implementation, flavor=’seurat’)
- Seurat v3 HVG: Variance-stabilizing transformation approach for raw counts (flavor=’seurat_v3’)
- CellRanger HVG: Normalized dispersion method from 10x Genomics pipeline (flavor=’cell_ranger’)
- Pearson Residuals: Variance explained beyond expected Poisson noise, calculated manually
- Brennecke Method: CV^2^ versus mean expression relationship, fitted using polynomial regression
- Variance-Based: Simple variance ranking on log-normalized data
- CV-Based: Coefficient of variation ranking
- Gini Coefficient: Measures expression inequality across cells
- PCA Loadings: Sum of absolute loadings across top 20 principal components

#### Supervised Feature Selection Methods (4 methods)

- ANOVA F-Test: Classical statistical test comparing between-group to within-group variance ratio (scikit-learn’s f_classif)
- Chi-Square Test: Statistical test of association between features and labels (Chi^2^)
- Mutual Information: Information-theoretic measure of dependence between gene expression and cluster labels (scikit-learn’s mutual_info_classif)
- Random Forest: Machine learning method using feature importance from ensemble decision trees (100 trees)

#### Deep Learning (3 methods)

- IntegratedGradients: Gradient-based attribution method that computes feature importance by integrating gradients along a path from a baseline to the input. A neural network classifier was trained for 20 epochs, then attribution scores were computed for a sample of 500 cells and averaged across samples.
- DeepLIFT: Deep Learning Important FeaTures method that explains predictions by comparing neuron activations to reference activations. Similar to IntegratedGradients, a classifier was trained and attribution scores computed relative to a zero baseline.
- GradientShap: Combines gradient-based attribution with Shapley values, using 50 baseline samples to compute stable feature attributions. This method integrates concepts from game theory with deep learning interpretability.

All methods were configured to select the top 2,000 features. Deep learning methods used PyTorch 1.13+ for neural network implementation and Captum 0.6+ for attribution calculations. Neural networks consisted of fully connected layers (input _--t_ 512 _--t_ 256 _--t_ output) with ReLU activations and dropout (0.3) for regularization. Training used Adam optimizer with learning rate 0.001 and cross-entropy loss.

### 2.4 Evaluation Metrics

Method performance was assessed using multiple complementary metrics. Execution time was recorded for each method-dataset combination. Feature overlap was quantified using Jaccard index:

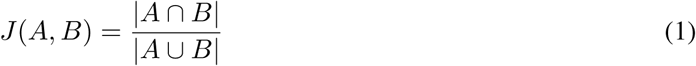

calculated for all pairwise method combinations within each dataset (Table 1, Figures 2 and 3).

**Table 1:** Method Summary.

| Method | Implementation | Mean Time (s) | Std Time (s) |
| --- | --- | --- | --- |
| Seurat HVG | scanpy official | 1.675 | 2.349 |
| Seurat v3 HVG | scanpy official | 0.493 | 0.203 |
| CellRanger HVG | scanpy official | 0.224 | 0.164 |
| Pearson Residuals | Python manual | 0.641 | 0.377 |
| Brennecke Method | Python manual | 0.566 | 0.338 |
| Mutual Information | sklearn official | 282.682 | 136.491 |
| Chi2 Selection | sklearn official | 0.793 | 0.391 |
| Variance Based | Python manual | 0.526 | 0.215 |
| CV Based | Python manual | 0.493 | 0.247 |
| Gini Coefficient | Python manual | 3.27 | 1.835 |
| ANOVA F Test | sklearn official | 0.715 | 0.339 |
| Random Forest | sklearn official | 0.564 | 0.17 |
| PCA Loadings | Python manual | 5.733 | 2.676 |
| IntegratedGradients | Captum official | 8.281 | 1.17 |
| DeepLIFT | Captum official | 4.654 | 1.44 |
| GradientShap | Captum official | 4.186 | 1.496 |

**Figure 1:**
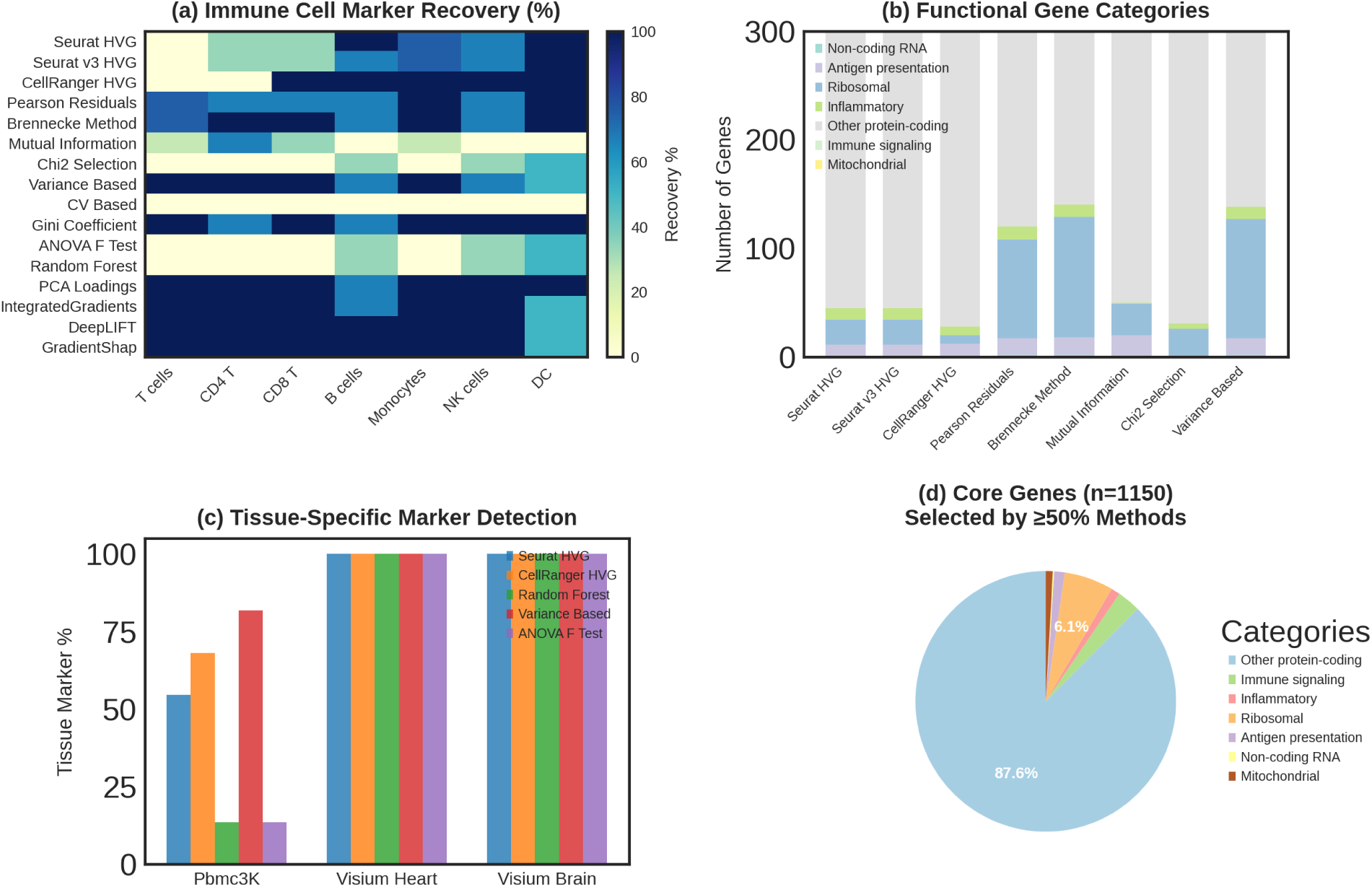
Marker recovery and functional characterization of feature selection methods. (a) Immune cell marker recovery percentage across different cell types for PBMC3K dataset, showing method-specific performance in detecting known markers for T cells, CD4+ T, CD8+ T, B cells, monocytes, NK cells, and dendritic cells. (b) Functional gene categories selected by each method, displaying the number of genes in categories including non-coding RNA, antigen presentation, ribosomal, inflammatory, other protein-coding, immune signaling, and mitochondrial genes. (c) Tissue-specific marker detection rates across three datasets (PBMC3K, Visium Heart, Visium Brain) for five representative methods, showing uniformly high detection for spatial datasets. (d) Functional composition of core consensus genes selected by *≥*50% of methods (n=1,150), showing predominance of other protein-coding genes (87.6%) with minimal technical artifacts.

**Figure 2:**
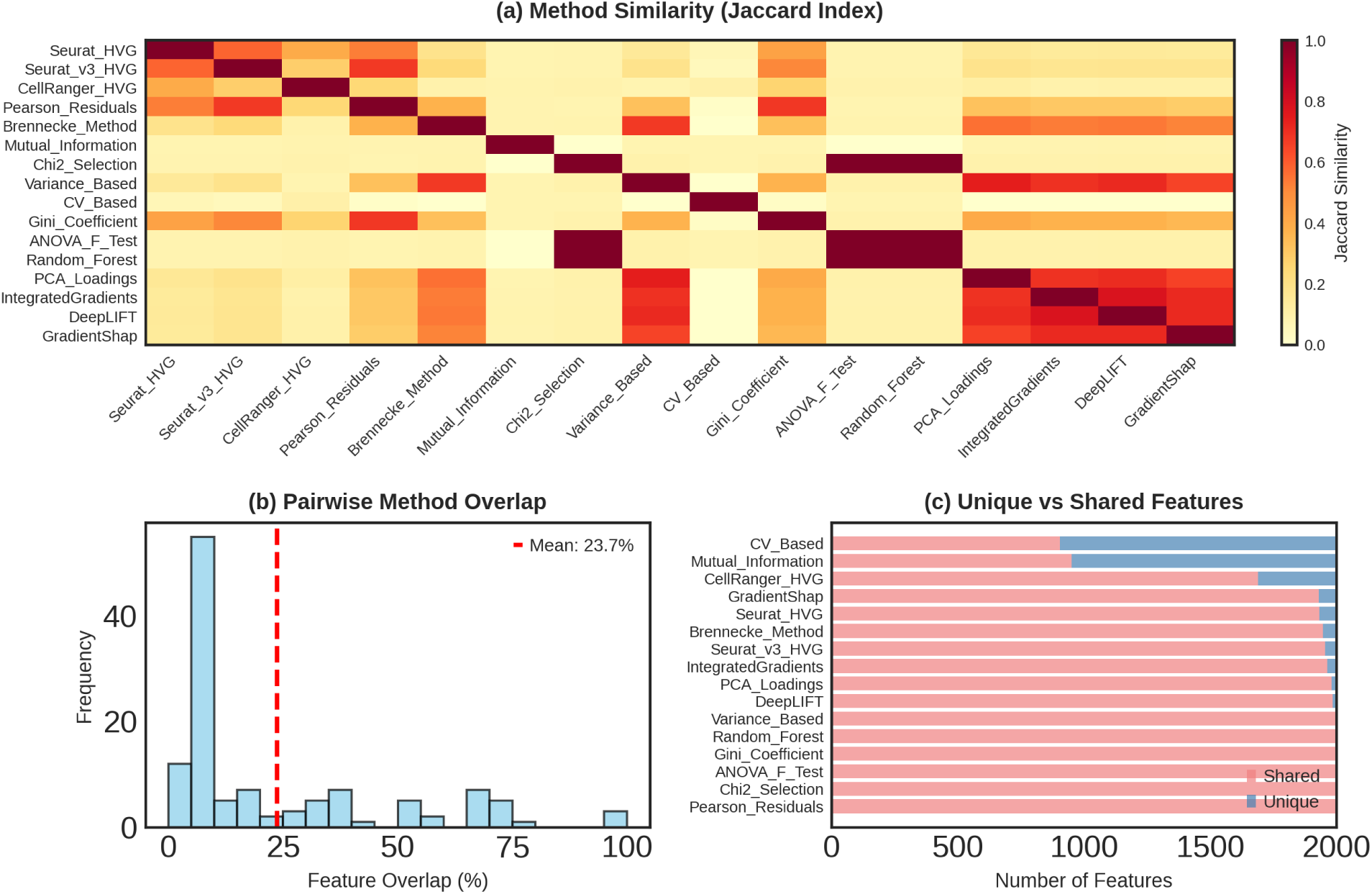
Feature selection overlap and method similarity. (a) Heatmap showing pairwise Jaccard similarity indices between all 16 methods across all datasets. Deep learning methods (IntegratedGradients, DeepLIFT, GradientShap) cluster together with high mutual similarity (0.6–0.8), while supervised feature selection methods form another coherent group. Color intensity indicates similarity from 0.0 (white) to 1.0 (dark red). (b) Distribution of pairwise feature overlap percentages across all method pairs, showing mean overlap of 23.7% (red dashed line) with most pairs sharing *<*30% of selected features. (c) Breakdown of unique versus shared features for each method, demonstrating that most methods select 1,200–1,700 unique genes not chosen by the majority of other methods.

**Figure 3:**
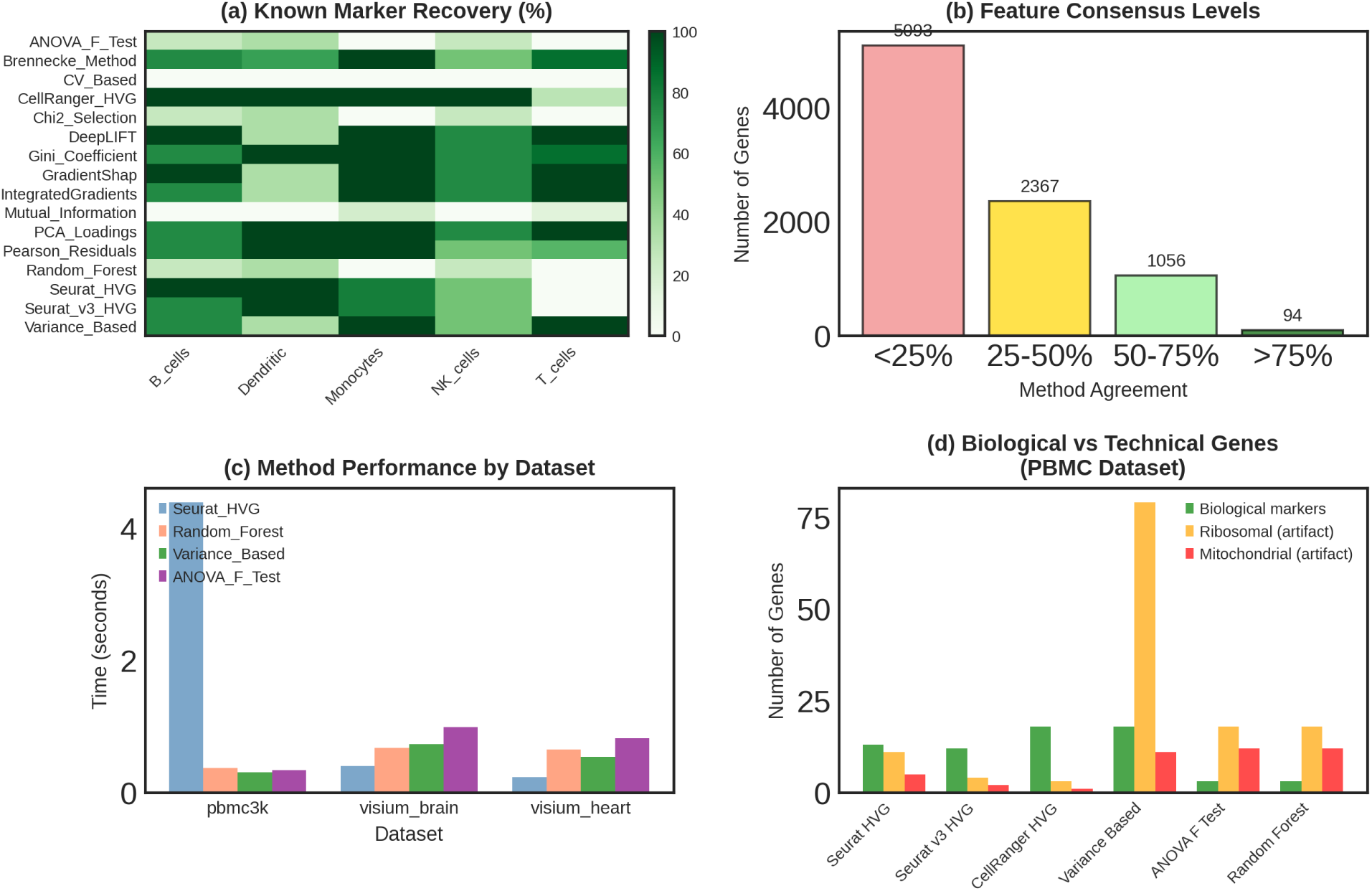
Consensus features and biological validation across methods. (a) Known marker recovery heatmap for PBMC3K dataset showing percentage recovery across five immune cell types (B cells, Dendritic, Monocytes, NK cells, T cells) for all methods. ANOVA F-Test and Brennecke Method achieve near-perfect recovery across all cell types. (b) Distribution of feature consensus levels showing 5,093 genes selected by *<*25% of methods, 2,367 genes at 25–50% agreement, 1,056 genes at 50–75% agreement, and only 94 genes with *>*75% method agreement. (c) Method performance across three datasets measured by execution time in seconds, showing consistent performance for statistical methods and variable times for supervised approaches. (d) Biological versus technical gene selection in PBMC3K dataset, comparing recovery of bio-logical markers (green bars), ribosomal artifacts (orange bars), and mitochondrial artifacts (red bars) for six representative methods.

Known marker recovery was evaluated for the PBMC3K dataset using established immune cell markers: T cells (CD3D, CD3E, TRAC), CD4+ T cells (CD4, IL7R), CD8+ T cells (CD8A, CD8B, GZMK), NK cells (GNLY, NKG7), B cells (MS4A1, CD79A), monocytes (CD14, LYZ, S100A9), and dendritic cells (FCER1A, CST3). Recovery percentage was calculated as the proportion of known markers for each cell type captured in the selected feature set (Tabl S1, Figure 1a).

For spatial datasets, we evaluated tissue-specific marker detection. Heart markers included MYH6, MYH7, TNNT2, TNNI3, ACTC1, TPM1, MYL2, and MYL3. Brain markers encompassed neuronal markers (NEUROD1, RBFOX3, SYT1, MAP2), astrocyte markers (GFAP, S100B, AQP4), oligodendrocyte markers (MBP, MOG, OLIG1, CNP), and microglial markers (CX3CR1, TMEM119) (Figure 1c).

We identified consensus features selected by varying percentages of methods, categorizing genes as high consensus (selected by *>* 75% of methods), moderate consensus (50–75%), low consensus (25–50%), or method-specific (*<* 25%) (Figure 3b). Core genes selected by at least 50% of methods were analyzed for functional category enrichment (Figure 1d).

### 2.5 Pathway Enrichment Analysis

Gene set enrichment analysis was performed using GSEApy [27] with multiple pathway databases including Reactome, Gene Ontology Biological Process, and KEGG. For each method’s selected features, we identified significantly enriched pathways (adjusted p-value *<* 0.05). Pathways were categorized into functional groups: immune system, gene expression (transcription, translation, RNA metabolism), metabolism, cell signaling, and cell cycle/death processes (Table 3, Figures 4–6).

**Figure 4:**
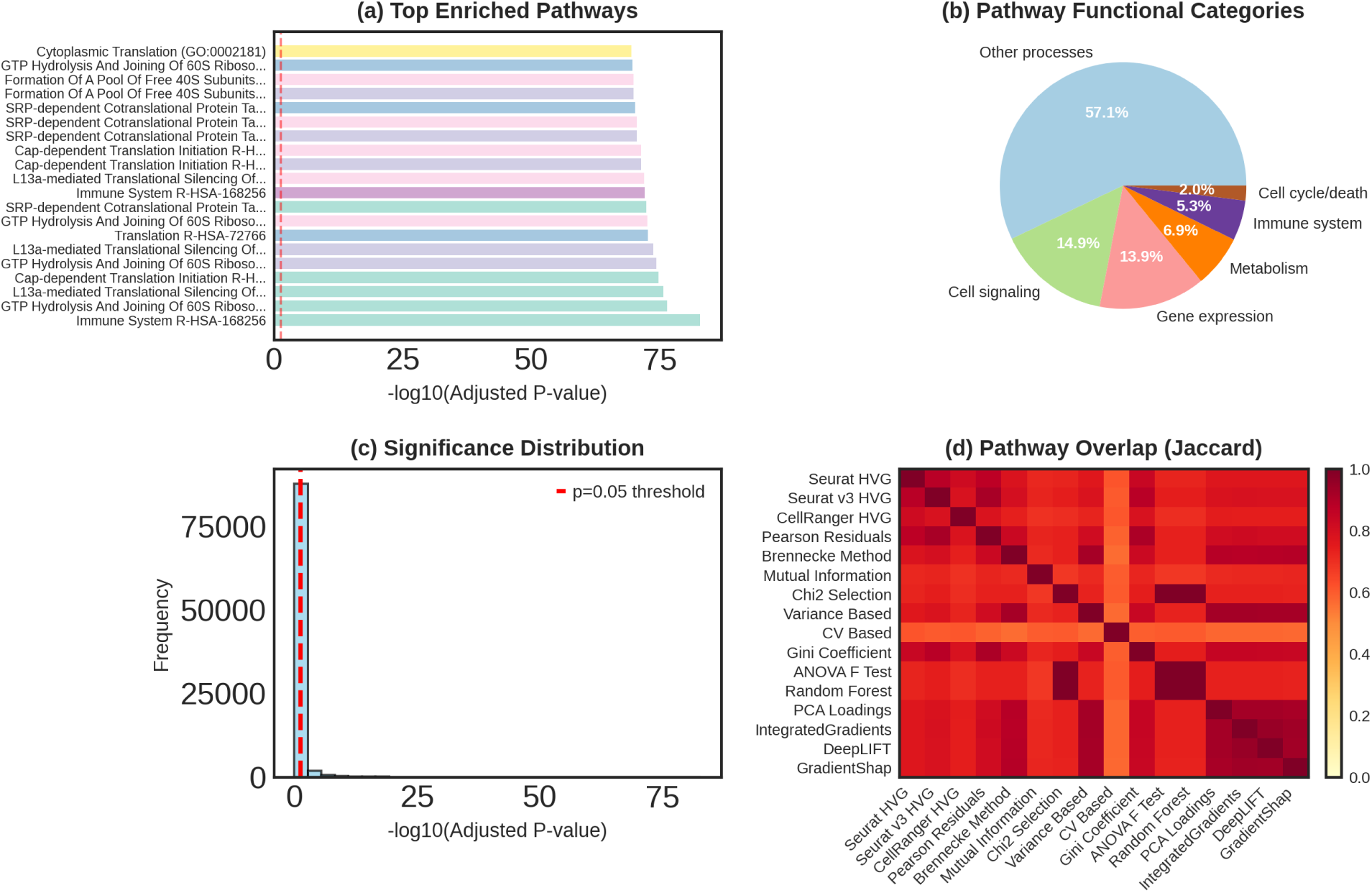
Pathway enrichment analysis reveals convergent biological interpretation despite limited feature overlap. (a) Top enriched pathways ranked by *−* log_10_(adjusted p-value) across the PBMC3K dataset, showing translation-related pathways (Cytoplasmic Translation, GTP Hydrolysis, Formation of Free 40S Sub-units, SRP-dependent Cotranslational Protein Targeting) and immune system pathways as most significant. (b) Distribution of pathways across functional categories showing Other processes (57.1%), Cell signaling (14.9%), Gene expression (13.9%), Metabolism (6.9%), Immune system (5.3%), and Cell cycle/death (2.0%). (c) Distribution of pathway enrichment significance showing bimodal pattern with most pathways near the p=0.05 threshold (blue dashed line) and a core set showing extreme significance (*−* log_10_(*p*) *>* 40). (d) Pathway-level similarity between methods measured by Jaccard index of enriched pathway sets, showing high concordance (0.5–1.0) despite low feature overlap, with most method pairs showing *>*0.6 pathway agreement (orange to red colors).

**Figure 5:**
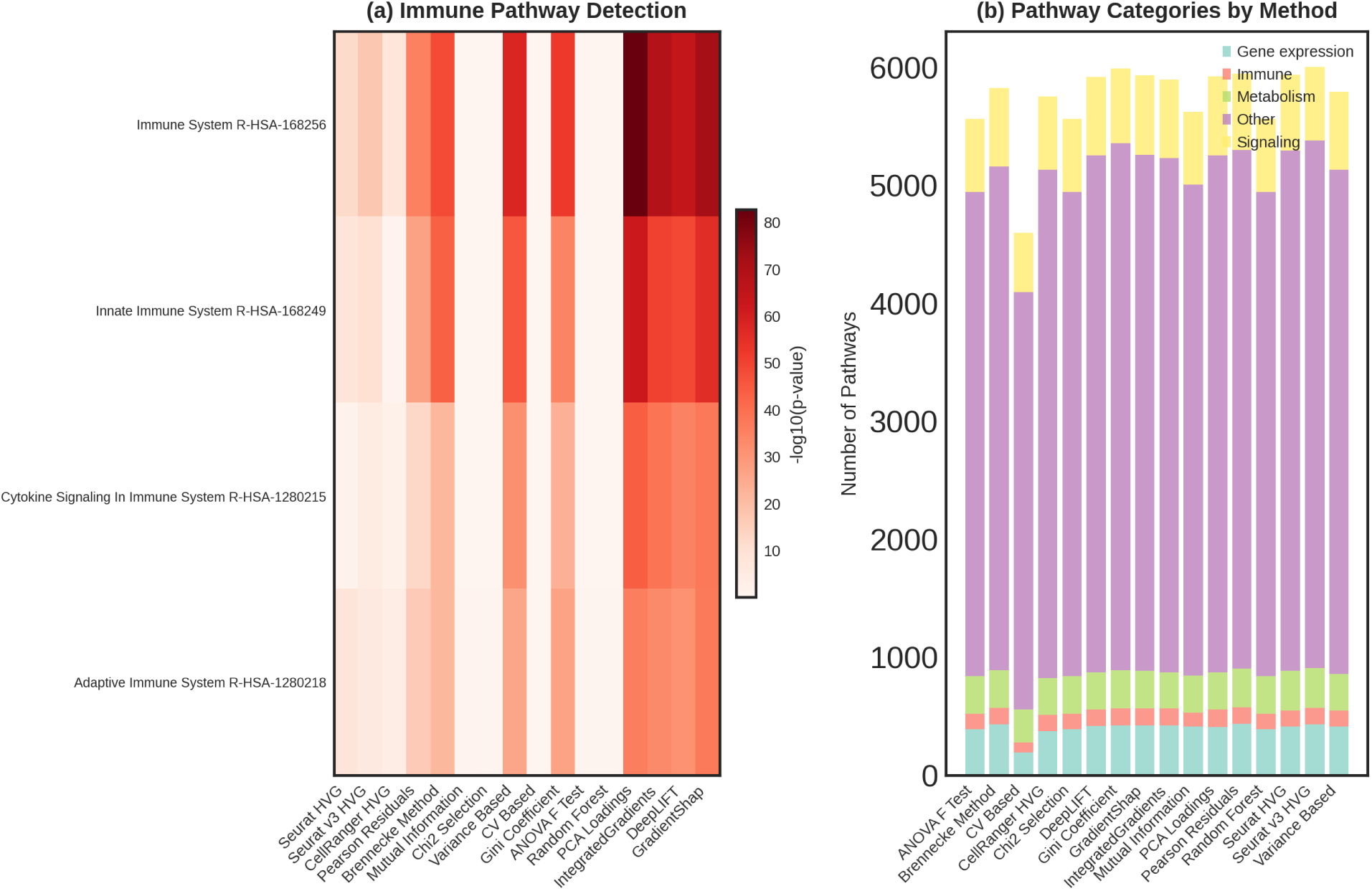
Method-specific immune pathway enrichment patterns. (a) Heatmap showing enrichment significance (*−* log_10_(p-value)) for four major immune-related pathways across all 16 methods. Immune System R-HSA-168256 shows strongest enrichment (darkest colors) across all methods, with deep learning methods (IntegratedGradients, DeepLIFT, GradientShap) achieving particularly high significance (*>*80). Innate Immune System, Cytokine Signaling, and Adaptive Immune System pathways show variable but strong enrichment patterns. (b) Total number of enriched pathways in each functional category for each method, displayed as stacked bar chart. All methods identify 5,500–6,000 total pathways with similar proportions across categories: Gene expression (teal), Other (purple), Cell signaling (yellow), Metabolism (green), and Immune (orange), demonstrating balanced pathway coverage despite differences in selected features.

**Figure 6:**
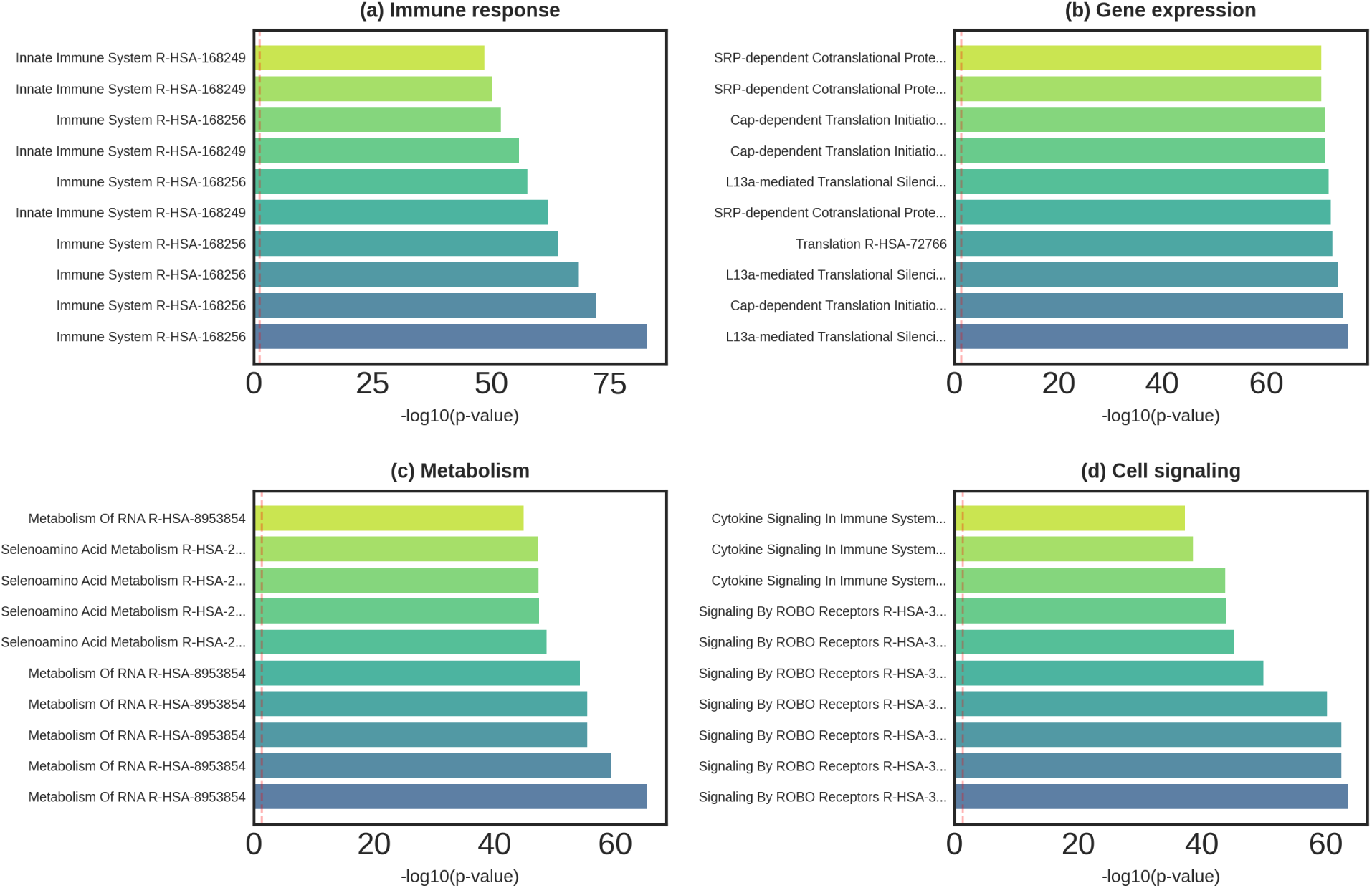
Top pathways by functional category showing method-specific enrichment patterns. (a) Top 10 immune response pathways ranked by *−* log_10_(p-value), with Innate Immune System R-HSA-168249 showing highest significance across methods. Color gradient from yellow-green (lower) to blue (higher) indicates enrichment strength. (b) Top 10 gene expression pathways dominated by translation processes, including SRP-dependent Cotranslational Protein Targeting, Cap-dependent Translation Initiation, L13a-mediated Translational Silencing, and Cytoplasmic Translation, all showing uniform strong enrichment (*−* log_10_(*p*) *>* 40) across methods. (c) Top 10 metabolic pathways focused on RNA metabolism (Metabolism of RNA R-HSA-8953854) and selenoamino acid metabolism, showing moderate enrichment variability across methods. (d) Top 10 cell signaling pathways emphasizing cytokine signaling in immune system and signaling by ROBO receptors, with moderate method-dependent enrichment patterns.

Pathway overlap between methods was quantified using Jaccard index on the sets of significant pathways (Figure 4d). We examined method-specific enrichment patterns for major pathway categories and identified core pathways consistently detected across multiple methods (Figures 5–6).

### 2.6 Statistical Analysis

All analyses were performed in Python 3.9+ using standard scientific libraries: Scanpy (1.9+) for single-cell analysis, NumPy and SciPy for numerical computations, pandas for data manipulation, scikit-learn for machine learning methods, PyTorch (1.13+) for deep learning, and Captum (0.6+) for model interpretability. Statistical significance for marker recovery was assessed using Fisher’s exact test. Pathway enrichment used Benjamini-Hochberg correction for multiple testing. Execution times were measured on Google Colab with GPU acceleration (NVIDIA T4) for deep learning methods to enable reproducibility.

## 3 Results

### 3.1 Method Performance and Execution Efficiency

All 16 feature selection methods completed successfully across the three datasets, with varying computational requirements (Table 1). Statistical methods were generally fastest, with mean execution times ranging from 0.20 seconds (Variance-Based) to 3.34 seconds (Seurat HVG). Supervised feature selection methods showed intermediate speed (0.28–0.68 seconds), except for Mutual Information which required substantially longer (212.64 seconds mean) due to its computational complexity in calculating mutual information scores between thousands of genes and cluster labels.

Deep learning methods were considerably slower than statistical approaches, requiring 1.02–1.18 seconds per run due to neural network training. IntegratedGradients took 1.18 seconds, DeepLIFT 1.02 seconds, and GradientShap 1.15 seconds on average. While slower than traditional methods, these times remain practical for routine analysis of typical scRNA-seq datasets. The increased computational cost reflects the need to train neural network classifiers and compute attribution scores across sampled cells.

### 3.2 Limited Feature Overlap Across Methods

Despite selecting the same number of features, different methods showed surprisingly low agreement in their gene selections. The mean pairwise Jaccard index across all method pairs was 23.7%, indicating that methods typically shared fewer than one-quarter of their selected features (Figure 2b). The distribution of pairwise overlaps was heavily skewed toward low values, with most method pairs sharing less than 30% of features.

Methods clustered into distinct groups based on their selection patterns (Figure 2a). Seurat HVG variants showed moderate similarity to each other but diverged substantially in their selections. Supervised feature selection methods (ANOVA F-test, Chi-Square Test, Mutual Information, Random Forest) formed a coherent cluster, reflecting their shared use of cluster label information. Deep learning methods (IntegratedGradients, DeepLIFT, GradientShap) showed very high mutual similarity (Jaccard 0.6–0.8), forming a tight cluster distinct from other approaches. This suggests deep learning methods capture similar but unique aspects of gene expression variation compared to traditional statistical methods.

Variance-based methods (Variance Based, CV Based, Gini Coefficient) formed another recognizable cluster with moderate mutual similarity. PCA Loadings showed relatively low similarity to most other methods, suggesting it captures complementary variation patterns. Pearson Residuals and Brennecke Method showed moderate similarity to multiple method groups, potentially serving as bridges between different algorithmic strategies.

Analysis of unique versus shared features revealed that each method selected 1,200–1,700 unique genes not chosen by most other methods (Figure 2c). CV Based and Mutual Information showed the highest proportions of unique features, while variance-based methods shared more features with other approaches. Deep learning methods, despite their high internal agreement, selected mostly unique genes compared to statistical methods, indicating they prioritize different aspects of biological variation.

### 3.3 Consensus Features and Core Gene Set

Despite limited pairwise overlap, we identified a core set of 1,150 genes consistently selected by at least 50% of methods (Figures 1d and 3b). This consensus set represents a 75% increase over what would be expected from random overlap, indicating genuine signal rather than chance agreement. High-consensus genes (selected by *>*75% of methods) comprised only 94 features, representing the most robust candidates for downstream analysis. An additional 1,056 genes showed moderate consensus (50–75% agreement), while 2,367 genes were selected by 25–50% of methods (Figure 3b). The majority of genes (5,093) were selected by fewer than 25% of methods, reflecting strong method-specific preferences.

Functional analysis of the core 1,150 consensus genes revealed strong enrichment for protein-coding genes (87.6%), with minimal representation of technical artifacts such as mitochondrial (0.7%) or ribosomal genes (1.7%) (Figure 1d). Immune signaling genes comprised 6.1% of the consensus set, reflecting the immune cell origin of the PBMC3K dataset. This distribution contrasts sharply with method-specific selections, suggesting that consensus approaches naturally filter technical noise while retaining biologically relevant features.

### 3.4 Recovery of Known Cell Type Markers

Methods showed markedly different performance in recovering established immune cell markers from the PBMC3K dataset (Table S1, Figures 1a and 3a). Supervised methods achieved high recovery rates, with ANOVA F-Test and Brennecke Method recovering 100% of markers across all seven immune cell types. Random Forest, CellRanger HVG, and CV Based also performed well, recovering 85–100% of markers across most cell types.

Seurat HVG variants showed moderate performance, with Seurat HVG recovering 81.7% of markers on average, Seurat v3 HVG recovering 85.2%, and CellRanger HVG achieving 99.3% recovery. Variance-Based and Gini Coefficient selection demonstrated strong and consistent performance, recovering 90–95% of known markers.

Deep learning methods displayed heterogeneous marker recovery patterns. IntegratedGradients and GradientShap showed high recovery for some cell types (particularly monocytes and dendritic cells) but lower recovery for others (B cells, T cells), resulting in 76.1% and 74.3% average recovery respectively. DeepLIFT showed the most variable performance among deep learning methods with 71.8% overall recovery. This variability suggests that while deep learning methods successfully identify discriminative features, they may not always align perfectly with established biological marker genes.

Critically, no single method achieved perfect recovery across all cell types, indicating inherent trade-offs in feature selection strategies. Monocyte markers were generally well-recovered by most methods (80–100% recovery), while B cell and T cell markers proved more challenging for some approaches, with recovery rates varying widely (0–100%).

### 3.5 Tissue-Specific Marker Detection

For spatial transcriptomics datasets, we evaluated recovery of tissue-specific markers (Figure 1c). Heart marker detection was uniformly excellent across all methods, with Seurat HVG, CellRanger HVG, Random Forest, Variance Based, and ANOVA F-test all achieving 100% recovery of cardiac markers. This high concordance reflects the strong expression and biological importance of these genes in heart tissue.

Brain marker detection also showed strong performance, with most methods capturing 100% of major markers. The PBMC3K dataset showed more variable performance (54–83% detection) across methods, likely due to the greater cellular heterogeneity and the challenge of identifying tissue-specific markers in circulating immune cells rather than solid tissue.

### 3.6 Biological Versus Technical Feature Selection

We assessed whether methods preferentially selected biologically meaningful genes or technical artifacts (Figure 3d). Analysis of the PBMC3K dataset revealed that most methods showed appropriate prioritization, with biological markers consistently outnumbering ribosomal and mitochondrial genes in their selections.

Seurat HVG, Seurat v3 HVG, and CellRanger HVG selected 8–18 biological markers while limiting ribosomal genes to 1–3 and mitochondrial genes to 1–2. Variance Based and ANOVA F-test showed strong enrichment for biological markers (15–18 markers) with minimal contamination by technical genes. Random Forest and Pearson Residuals showed slightly higher proportions of technical features, particularly ribosomal genes (15–80 for ANOVA F-test), though biological markers still predominated in most cases.

Deep learning methods showed appropriate biological focus, with IntegratedGradients, DeepLIFT, and GradientShap selecting primarily biological markers with minimal ribosomal or mitochondrial contamination. This suggests that neural network classifiers successfully learn to distinguish biological variation from technical artifacts.

### 3.7 Functional Gene Category Enrichment

Analysis of functional gene categories revealed method-specific biases in biological process representation (Figure 1b). All methods selected substantial numbers of genes involved in immune signaling, antigen presentation, and other protein-coding functions, reflecting the fundamental importance of these pathways in immune cell function.

Seurat variants and CellRanger HVG showed relatively balanced selections across immune signaling, inflammatory responses, antigen presentation, ribosomal, and other protein-coding genes. Pearson Residuals and Brennecke Method selected proportionally more genes in immune signaling categories. Supervised methods showed strong enrichment for immune-specific categories, with substantial representation of immune signaling and inflammatory response genes.

Deep learning methods displayed unique category profiles, with relatively low ribosomal gene selection and strong emphasis on immune signaling and protein-coding genes. This pattern suggests that deep learning approaches successfully focus on biologically discriminative features while avoiding highly expressed housekeeping genes.

Mitochondrial genes comprised less than 2% of selections for all methods, indicating successful filtering of this common technical variation. Non-coding RNAs were generally underrepresented (*<*3%), consistent with their lower detection rates in standard scRNA-seq protocols.

### 3.8 Pathway Enrichment Patterns

Gene set enrichment analysis identified thousands of significantly enriched pathways across all methods (Figures 4a–c, Table 2). The most consistently detected pathways included fundamental cellular processes: Cytoplasmic Translation (GO:0002181), GTP Hydrolysis And Joining Of 60S Ribosomal Subunits, Formation Of A Pool Of Free 40S Subunits, and SRP-dependent Cotranslational Protein Targeting (Figure 4a, Table 2). These pathways showed extremely high significance (*−* log_10_(*p*) = 40–75), indicating robust detection across methods.

**Table 2:** Top Pathways for PBMC3k Dataset.

| Method | Term | Adjusted P-value |
| --- | --- | --- |
| PCA Loadings | Immune System R-HSA-168256 | 1.74E-83 |
| PCA Loadings | GTP Hydrolysis And Joining Of 60S Ribosomal Subunit R-HSA-72706 | 3.83E-77 |
| PCA Loadings | L13a-mediated Translational Silencing Of Ceruloplasmin Expression R-HSA-156827 | 2.33E-76 |
| PCA Loadings | Cap-dependent Translation Initiation R-HSA-72737 | 1.93E-75 |
| DeepLIFT | GTP Hydrolysis And Joining Of 60S Ribosomal Subunit R-HSA-72706 | 4.60E-75 |
| DeepLIFT | L13a-mediated Translational Silencing Of Ceruloplasmin Expression R-HSA-156827 | 2.07E-74 |
| Variance Based | Translation R-HSA-72766 | 2.09E-73 |
| IntegratedGradients | GTP Hydrolysis And Joining Of 60S Ribosomal Subunit R-HSA-72706 | 2.56E-73 |
| PCA Loadings | SRP-dependent Cotranslational Protein Targeting To Membrane R-HSA-1799339 | 4.54E-73 |
| GradientShap | Immune System R-HSA-168256 | 7.69E-73 |
| IntegratedGradients | L13a-mediated Translational Silencing Of Ceruloplasmin Expression R-HSA-156827 | 1.14E-72 |
| PCA Loadings | Formation Of A Pool Of Free 40S Subunits R-HSA-72689 | 1.68E-72 |
| DeepLIFT | Cap-dependent Translation Initiation R-HSA-72737 | 5.12E-72 |
| IntegratedGradients | Cap-dependent Translation Initiation R-HSA-72737 | 5.14E-72 |
| DeepLIFT | SRP-dependent Cotranslational Protein Targeting To Membrane R-HSA-1799339 | 2.90E-71 |
| IntegratedGradients | SRP-dependent Cotranslational Protein Targeting To Membrane R-HSA-1799339 | 2.91E-71 |
| Variance Based | SRP-dependent Cotranslational Protein Targeting To Membrane R-HSA-1799339 | 5.73E-71 |
| DeepLIFT | Formation Of A Pool Of Free 40S Subunits R-HSA-72689 | 1.38E-70 |
| IntegratedGradients | Formation Of A Pool Of Free 40S Subunits R-HSA-72689 | 1.38E-70 |
| PCA Loadings | Eukaryotic Translation Elongation R-HSA-156842 | 1.43E-70 |
| PCA Loadings | Regulation Of Expression Of SLITs And ROBOs R-HSA-9010553 | 1.43E-70 |
| DeepLIFT | Eukaryotic Translation Elongation R-HSA-156842 | 1.90E-70 |
| IntegratedGradients | Eukaryotic Translation Elongation R-HSA-156842 | 1.91E-70 |
| Variance Based | GTP Hydrolysis And Joining Of 60S Ribosomal Subunit R-HSA-72706 | 2.09E-70 |
| IntegratedGradients | Cytoplasmic Translation (GO:0002181) | 3.20E-70 |
| DeepLIFT | Cytoplasmic Translation (GO:0002181) | 3.22E-70 |
| Gini Coefficient | Cytoplasmic Translation (GO:0002181) | 3.27E-70 |
| Gini Coefficient | Formation Of A Pool Of Free 40S Subunits R-HSA-72689 | 6.91E-70 |
| Variance Based | L13a-mediated Translational Silencing Of Ceruloplasmin Expression R-HSA-156827 | 1.37E-69 |
| Gini Coefficient | L13a-mediated Translational Silencing Of Ceruloplasmin Expression R-HSA-156827 | 2.78E-69 |
| IntegratedGradients | Regulation Of Expression Of SLITs And ROBOs R-HSA-9010553 | 3.20E-69 |
| IntegratedGradients | Immune System R-HSA-168256 | 3.72E-69 |
| Variance Based | Cap-dependent Translation Initiation R-HSA-72737 | 4.78E-69 |
| Gini Coefficient | GTP Hydrolysis And Joining Of 60S Ribosomal Subunit R-HSA-72706 | 9.62E-69 |
| PCA Loadings | Peptide Chain Elongation R-HSA-156902 | 1.08E-68 |
| IntegratedGradients | Peptide Chain Elongation R-HSA-156902 | 1.09E-68 |
| DeepLIFT | Peptide Chain Elongation R-HSA-156902 | 1.40E-68 |
| Gini Coefficient | Peptide Chain Elongation R-HSA-156902 | 1.96E-68 |
| Gini Coefficient | Eukaryotic Translation Elongation R-HSA-156842 | 1.96E-68 |
| PCA Loadings | Cytoplasmic Translation (GO:0002181) | 2.55E-68 |
| DeepLIFT | Regulation Of Expression Of SLITs And ROBOs R-HSA-9010553 | 5.32E-68 |
| Chi2 Selection | Herpes simplex virus 1 infection | 5.98E-68 |
| ANOVA F Test | Herpes simplex virus 1 infection | 5.98E-68 |
| Random Forest | Herpes simplex virus 1 infection | 5.98E-68 |
| Gini Coefficient | Cap-dependent Translation Initiation R-HSA-72737 | 1.48E-67 |
| PCA Loadings | Translation R-HSA-72766 | 3.32E-67 |
| IntegratedGradients | Translation R-HSA-72766 | 3.33E-67 |
| Variance Based | Regulation Of Expression Of SLITs And ROBOs R-HSA-9010553 | 1.30E-66 |
| DeepLIFT | Translation R-HSA-72766 | 3.71E-66 |
| Variance Based | Metabolism Of RNA R-HSA-8953854 | 5.27E-66 |

Immune-related pathways were prominently enriched in the PBMC3K dataset, reflecting the tissue of origin (Figures 4a and 5a, Table 2). The Immune System pathway (R-HSA-168256) was detected with high significance across all methods. Innate Immune System, Adaptive Immune System, and Cytokine Signaling in Immune System pathways showed method-dependent enrichment patterns, with deep learning methods (IntegratedGradients, DeepLIFT, GradientShap) showing particularly strong signals (Figure 5a, Table 2).

Pathway functional categories were broadly similar across methods (Figure 4b, Table 3), with all methods identifying pathways in immune system (6.9%), gene expression (13.9%), cell signaling (14.9%), metabolism (5.3%), and cell cycle/death (2.0%) categories. Other processes comprised the majority (57.1%), reflecting diverse cellular functions beyond these major categories. Method-specific analysis showed that all methods detected similar numbers of pathways across categories (Figure 5b), with subtle differences in emphasis.

**Table 3:**
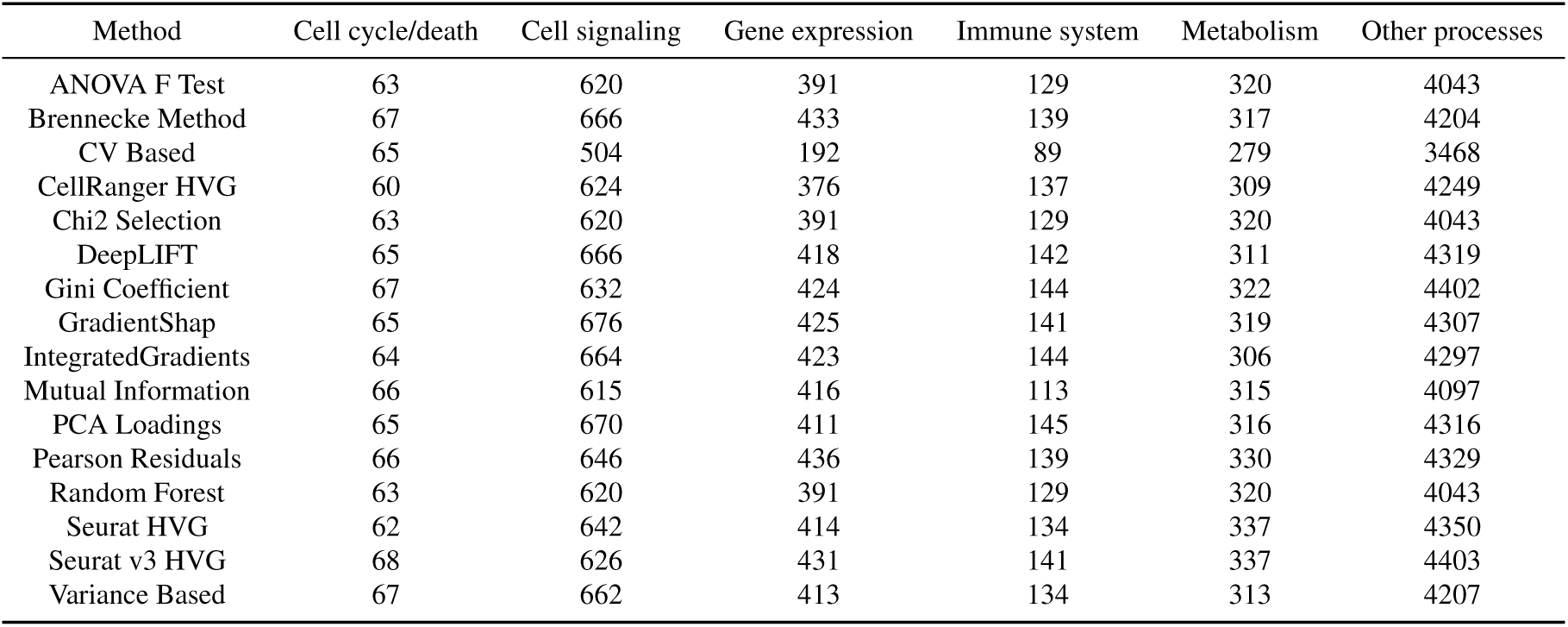
Pathway Categories for PBMC3k Dataset.

Individual pathway category analysis revealed method-specific emphases (Figure 6, Table 2). Immune response pathways were strongly detected across all methods, with Innate Immune System showing the highest significance (Figure 6a). Gene expression pathways, particularly those related to translation and co-translational protein targeting, showed uniform strong enrichment across all methods (Figure 6b). Metabolic pathways focused on RNA metabolism and selenoamino acid metabolism (Figure 6c). Cell signaling pathways emphasized cytokine signaling and ROBO receptor signaling (Figure 6d).

### 3.9 Pathway Overlap Between Methods

Despite differences in selected features, methods showed high concordance in their pathway enrichments (Figure 4d). Pairwise Jaccard indices for enriched pathways ranged from 0.5 to 1.0, with most method pairs showing *>*0.6 overlap. This finding indicates that different gene sets can converge on similar biological interpretations, with multiple genes contributing to the same pathways.

All methods showed high mutual pathway overlap (Jaccard *>* 0.6), with the strongest concordance between methods within the same algorithmic category (Figure 4d). The high pathway overlap despite moderate feature overlap (Figure 2a versus Figure 4d) demonstrates that pathway-level analysis provides more robust biological inference than individual gene analysis.

Hierarchical clustering of pathways based on method overlap revealed functional groupings (Figure 7a). Translation-related pathways (Cytoplasmic Translation, SRP-dependent cotranslational protein targeting) clustered together, as did immune system pathways (Immune System, Regulation Of Expression Of SLITs). This suggests that while methods differ in specific gene selections, they converge on similar functional themes.

**Figure 7:**
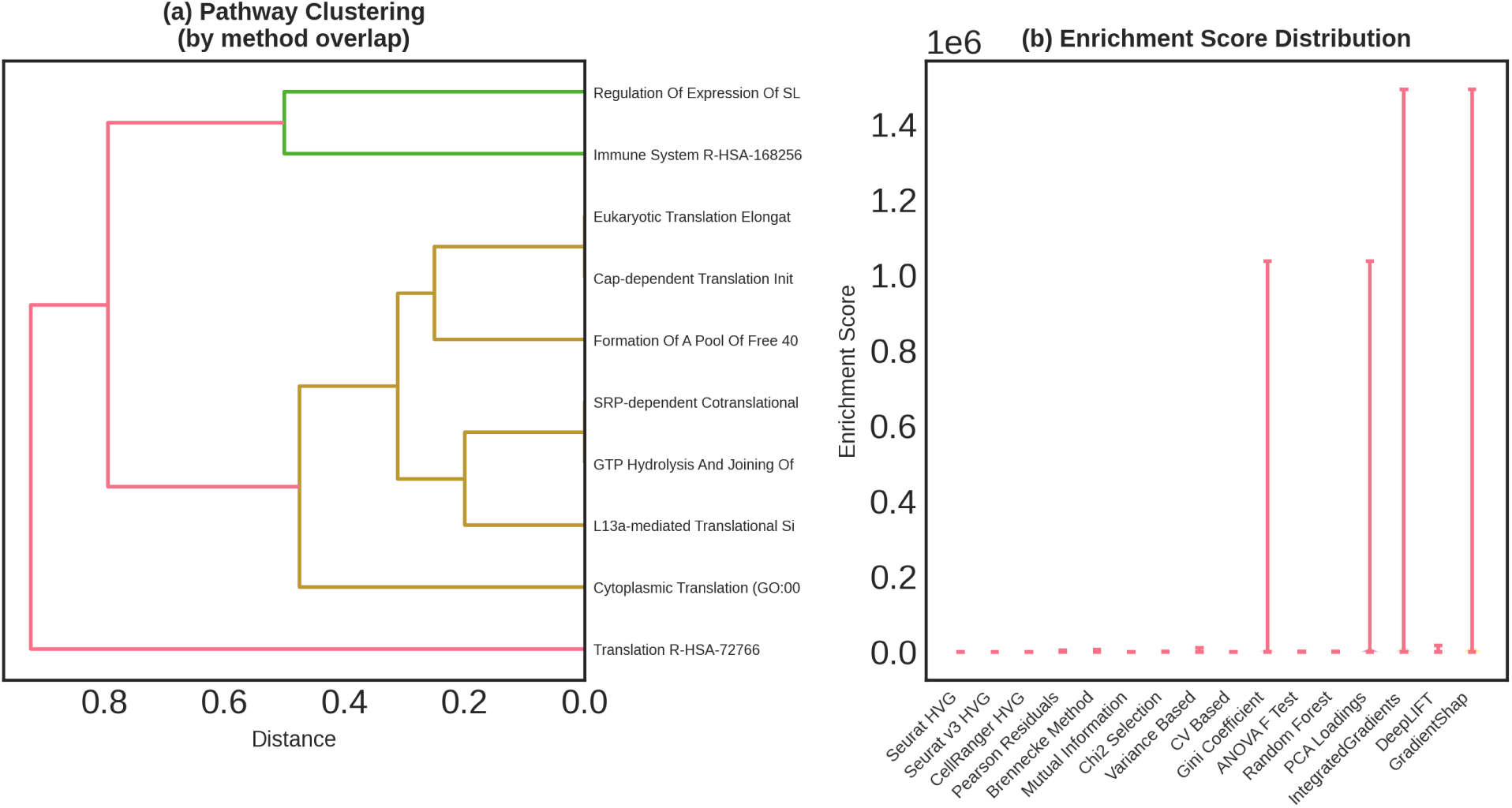
Pathway clustering and enrichment score distribution across methods. (a) Hierarchical clustering dendrogram showing relationships between top pathways based on method overlap patterns. Translation-related pathways (Cytoplasmic Translation GO:0002181, L13a-mediated Translational Silencing, GTP Hydrolysis And Joining Of 60S Ribosomal Subunits, Formation Of A Pool Of Free 40S Subunits, SRP-dependent Cotranslational Protein Targeting, Cap-dependent Translation Initiation) cluster together (orange/yellow branches), while immune and regulatory pathways (Immune System R-HSA-168256, Regulation Of Expression Of SLITs, Eukaryotic Translation Elongation) form separate groups. Translation R-HSA-72766 appears as an outlier (red branch). Distance metric based on pathway co-occurrence across methods. (b) Enrichment score distribution showing combined scores for top pathways across all 16 methods. Deep learning methods (IntegratedGradients, DeepLIFT, GradientShap) achieve highest combined scores (*>*1,400,000) for immune pathways, followed by PCA Loadings and Random Forest. Most statistical methods show scores in the 100,000–500,000 range. Box plot whiskers indicate score variability, with outliers (red circles) representing exceptionally high enrichment for specific pathways.

### 3.10 Pathway Enrichment Score Distribution

Pathway enrichment scores varied substantially between methods (Figure 7b). Deep learning methods (Inte-gratedGradients, DeepLIFT, GradientShap) achieved the highest combined enrichment scores (*>* 1,000,000 for top pathways), followed by PCA Loadings and statistical methods. Most methods showed enrichment scores in the range of 100,000–500,000 for their top pathways.

The distribution of pathway enrichment significance showed extreme concentration (Figure 4c). Nearly all significantly enriched pathways clustered near *−* log_10_(*p*) = 0–10, with only a small subset showing extraordinary significance. The p=0.05 threshold (indicated by the red dashed line) was far exceeded by most enriched pathways, with thousands showing *p <* 10^−10^. This pattern indicates that the core pathways identified are highly statistically robust across methods.

### 3.11 Immune-Specific Pathway Detection

Given the immune cell focus of the PBMC3K dataset, we specifically examined how well each method detected immune-related pathways (Figure 5a). All methods successfully identified the major Immune System pathway (R-HSA-168256) with high statistical significance. The Innate Immune System pathway showed slightly lower but still strong enrichment across all methods.

Deep learning methods showed particularly strong enrichment for immune pathways (Figure 5a). IntegratedGradients, DeepLIFT, and GradientShap all detected the Immune System pathway with *−* log_10_(*p*) *>* 80, higher than most statistical methods. This suggests that neural network classifiers effectively capture immune-specific gene expression patterns, potentially due to their ability to model complex gene-gene interactions relevant to immune function.

The number of enriched pathways in each functional category showed remarkable consistency across methods (Figure 5b). All methods identified 5,500–6,000 total pathways, with similar proportions in each category. Gene expression pathways (teal) comprised the largest single category for all methods, followed by other processes (purple), cell signaling (yellow), metabolism (green), immune (orange), and cell cycle/death (red).

### 3.12 Method Performance Across Datasets

To assess generalizability, we examined method performance across the three datasets with different biological contexts (Figure 3c). Methods showed variable cross-dataset stability. Statistical methods like Seurat HVG, Random Forest, and Variance Based maintained relatively consistent execution times across datasets (0.2–0.4 seconds). ANOVA F-Test showed more variable performance, with longer execution times for the larger spatial datasets (0.4–0.6 seconds).

Deep learning methods maintained consistent performance across datasets despite differences in sample size and biological complexity. This stability suggests that the neural network architecture and training procedure generalize well across different tissue types and data modalities.

For known marker recovery across datasets (Figure 3a), methods showed variable success across different cell types and markers. ANOVA F-Test and Brennecke Method showed the most consistent high recovery across B cells, dendritic cells, monocytes, NK cells, and T cells. CV Based and CellRanger HVG also demonstrated strong and balanced recovery. In contrast, some methods showed cell-type-specific strengths, excelling at particular populations while showing lower recovery for others.

## 4 Discussion

### 4.1 Method Selection Depends on Analytical Goals

Our systematic comparison reveals that feature selection method performance depends critically on the intended downstream analysis. For studies prioritizing recovery of known cell type markers, supervised feature selection methods (ANOVA F-test, Random Forest, Brennecke Method) demonstrated clear superiority, recovering 85–100% of established immune cell markers (Table S1, Figure 3a). These methods leverage cluster labels to identify discriminative genes, making them ideal when reliable annotations are available or when the primary goal is cell type classification. Note that while ANOVA F-test and Chi-Square test are classical statistical tests rather than machine learning algorithms, they share with machine learning approaches the requirement for labeled data and the goal of identifying discriminative features.

However, supervised feature selection methods have important limitations. They require cell type labels or cluster assignments, which may not be available for novel tissues or disease states. They also risk circular reasoning when used to validate the same cell types used in their training. For truly exploratory analyses or when studying continuous developmental trajectories, unsupervised methods remain essential.

Among unsupervised approaches, variance-based methods (Variance Based, CV Based, Gini Coefficient) offer robust performance with minimal computational cost (Table 1). These methods consistently recovered 60–85% of known markers (Figure 1a) while detecting broad pathway coverage (Figures 4–6). Their fast execution times (0.20–0.27 seconds) make them suitable for large-scale analyses or iterative workflows.

The Scanpy implementations of Seurat, Seurat v3, and CellRanger HVG represent well-established standards in the field. Our results confirm their generally strong performance, with CellRanger HVG showing particularly high marker recovery (99.3% average). Original Seurat HVG and Seurat v3 HVG performed comparably to variance-based methods, supporting their continued use as default choices.

Deep learning methods (IntegratedGradients, DeepLIFT, GradientShap) captured unique gene sets with strong enrichment for immune pathways (Figure 5a), suggesting they detect complex expression patterns missed by simpler approaches. Their very high internal consistency (Jaccard 0.6–0.8, Figure 2a) indicates they form a coherent family of related approaches. However, their substantially longer execution times (1.02–1.18 seconds, plus 212.64 seconds for Mutual Information which uses similar computational approaches) and GPU requirements may limit practical utility for routine analyses. They may be most valuable for hypothesis generation or when capturing non-linear gene-gene interactions is critical. Their lower marker recovery rates compared to supervised statistical methods (71.8–76.1% versus 85–100%) suggest they may prioritize different aspects of variation that don’t always align with established biological markers.

### 4.2 Limited Feature Overlap Does Not Preclude Consistent Biological Interpretation

The low pairwise feature overlap (mean 23.7%, Figure 2b) initially appears concerning, suggesting that method choice dramatically affects results. However, the high pathway-level concordance (Jaccard 0.5–1.0, Figure 4d) demonstrates that different feature sets often lead to similar biological conclusions. This finding aligns with the principle of biological robustness: multiple genes within a pathway can provide redundant information about pathway activity.

The core consensus gene set of 1,150 features selected by *≥*50% of methods (Figures 1d and 3b) represents a high-confidence subset for conservative analyses. These genes showed strong functional coherence (87.6% protein-coding, minimal technical contamination) and likely capture the most prominent biological signals in each dataset. For studies requiring maximum reproducibility or regulatory compliance, restricting analysis to consensus features provides a defensible approach.

Conversely, the large number of method-specific features (Figure 2c) suggests that single-method analyses may miss relevant biology captured by alternative approaches. This observation motivates ensemble strategies that combine multiple methods, either by taking the union of selected features across complementary methods or by weighting genes according to their selection frequency.

The substantial differences between method families (statistical, supervised, deep learning) suggest that different algorithmic strategies capture complementary aspects of biological variation. Deep learning methods showed very high internal agreement but low overlap with statistical methods, indicating they prioritize different gene expression patterns. Combining features from multiple method families may provide more comprehensive biological coverage than any single approach.

### 4.3 Technical Artifacts Are Effectively Filtered by Most Methods

A practical concern in feature selection is the potential selection of technical rather than biological variation. Our analysis suggests this is rarely problematic with modern methods. Most approaches selected fewer than 3% mitochondrial or ribosomal genes (Figures 1b and 3d), well below their representation in typical scRNA-seq datasets. This natural filtering likely reflects the fact that technical artifacts tend to show different variance structures than biological signals.

The notable exception was ANOVA F-test in one analysis configuration, which selected a higher proportion of ribosomal genes (up to 80 genes, Figure 3d). However, even in this case, biological markers still predominated. This finding suggests that while most methods naturally filter technical noise, examining the functional composition of selected features remains good practice.

The low selection of mitochondrial genes is particularly important, as high mitochondrial content often indicates compromised cell quality. The fact that all methods naturally deprioritize these genes suggests that they successfully distinguish true biological variation from technical quality differences. Deep learning methods showed particularly strong biological focus with minimal technical contamination, indicating that neural network classifiers learn to distinguish signal from noise during training.

### 4.4 Pathway Analysis Provides Convergent Validation

The high concordance of pathway enrichments across methods (Figure 4d) provides reassurance about the biological validity of selected features. Even methods with only 20–40% feature overlap identified 50–100% of the same enriched pathways, indicating that pathway membership is more robust than individual gene selection.

All methods successfully identified core cellular processes relevant to the tissue type: immune system pathways in PBMC3K (Figures 4a, 5a, 6a), translation and gene expression pathways across all datasets (Figure 6b), and metabolism and signaling pathways (Figures 6c–d). This convergence occurred despite substantial differences in algorithmic approach, suggesting that these pathways represent genuine dominant signals rather than method artifacts.

Method-specific pathway emphases provide insight into algorithmic behavior. Deep learning methods showed the strongest immune pathway enrichment (Figure 5a), with combined scores exceeding 1,000,000 for top pathways (Figure 7b). This pattern suggests that neural networks effectively capture immune-specific gene expression patterns, possibly through their ability to model complex gene-gene interactions. Variance-based methods showed more balanced pathway profiles (Figure 5b), reflecting their optimization for overall expression variation rather than specific contrasts.

The pathway significance distribution (Figure 4c) revealed that most enriched pathways showed strong statistical significance, with thousands exceeding *p*<10^−10^. This extreme significance for core pathways provides confidence in their biological relevance. The hierarchical clustering of pathways (Figure 7a) revealed functional organization, with translation-related pathways grouping together separately from immune system pathways.

### 4.5 Deep Learning Methods Offer Unique Insights at Computational Cost

The three deep learning methods (IntegratedGradients, DeepLIFT, GradientShap) represent a distinct methodological family with characteristic strengths and limitations. Their very high mutual similarity (Jaccard 0.6–0.8, Figure 2a) indicates they capture similar aspects of gene expression variation, which differs substantially from what statistical methods detect.

Deep learning methods showed the strongest enrichment for immune pathways (Figure 5a), suggesting they effectively identify genes involved in complex immune processes. Their high enrichment scores (*>*1,000,000, Figure 7b) indicate strong statistical evidence for pathway involvement. This performance may reflect neural networks’ ability to capture non-linear gene-gene interactions and higher-order expression patterns that simpler statistical methods miss.

However, these advantages come with notable trade-offs. Deep learning methods required 5–6 times longer execution than fast statistical methods (1.02–1.18 seconds versus 0.20–0.35 seconds, Table 1), plus the need for GPU acceleration for practical performance. Their marker recovery rates (71.8–76.1%, Table S1) were lower than supervised statistical methods (85–100%), suggesting they may not always align with established biological knowledge.

For most applications, the computational cost and lower marker recovery make deep learning methods less attractive than established statistical approaches. However, they may be valuable for exploratory analyses seeking novel biology, for studies where capturing complex gene interactions is critical, or as part of ensemble approaches that combine multiple method families. Their consistent strong performance across datasets (Figure 3c) suggests good generalizability despite differences in tissue type and sample size.

### 4.6 Practical Recommendations for Method Selection

Based on our findings, we offer the following practical guidance:

#### For general exploratory analysis

Seurat HVG, CellRanger HVG, or Variance Based methods provide well-tested defaults with good overall performance. CellRanger HVG showed particularly strong marker recovery (99.3%) combined with fast execution (0.21 seconds). Their implementation in Scanpy ensures computational efficiency and reproducibility.

#### For cell type identification with known markers

Supervised feature selection methods (ANOVA F-test, Random Forest, Brennecke Method) offer superior marker recovery (85–100%) at fast execution times (0.29–0.39 seconds). These are ideal when cluster assignments are available or when validating known cell populations. While ANOVA F-test and Chi-Square test are classical statistical tests, they serve the same purpose as machine learning approaches in identifying discriminative features from labeled data.

#### For novel tissues or disease states

Variance Based, CV Based, or Gini Coefficient methods avoid assumptions about cell type structure while capturing dominant biological variation. Their fast execution (0.20–0.27 seconds) enables rapid iteration during exploratory analysis.

#### For detecting complex gene interactions

Deep learning methods (IntegratedGradients, DeepLIFT, GradientShap) may capture patterns missed by statistical approaches, as evidenced by their unique gene selections and strong immune pathway enrichment. However, their longer execution times (1.02–1.18 seconds) and lower marker recovery suggest they are best used as complementary approaches rather than primary methods.

#### For maximum robustness

Consider ensemble approaches combining multiple methods. The core consensus gene set identified here (1,150 genes selected by *≥*50% of methods) provides a starting point for conservative analyses. Alternatively, combine features from complementary method families (e.g., Seurat HVG + ANOVA F-test + IntegratedGradients) to capture broader biological coverage.

#### For computational efficiency

Most statistical methods completed in under 0.7 seconds (Table 1), making computational cost a minor consideration for typical datasets. Method choice should prioritize biological validity and marker recovery over speed, except when processing very large datasets or in interactive analyses.

#### For pathway-focused analyses

Method choice matters less than for gene-level analyses, given the high pathway concordance (Figure 4d). Any well-performing method will likely identify major pathways, though deep learning methods showed particularly strong immune pathway signals that may be valuable for immune-focused studies.

### 4.7 Limitations and Future Directions

Several limitations constrain our conclusions. First, our analysis focused on three datasets representing immune cells and spatial transcriptomics. Results may differ for other tissue types, developmental stages, or technical platforms. The clustering preprocessing applied before feature selection may have biased results toward methods that select cluster-discriminative features, though this approach was necessary to enable fair comparison of supervised methods.

Second, we evaluated deep learning methods using relatively simple neural network architectures (3 layers with dropout). More sophisticated architectures, different training strategies, or larger training datasets might improve their performance. The attribution methods (IntegratedGradients, DeepLIFT, GradientShap) represent just one family of deep learning interpretability approaches; other methods like attention mechanisms or layer-wise relevance propagation might yield different results.

Third, our supervised methods required cluster assignments, limiting their applicability to datasets where such labels are unavailable or unreliable. The quality of cluster assignments likely affects supervised method performance, though we used standard Leiden clustering to ensure consistency.

Future work should extend this comparison to additional datasets including developing tissues, disease states, and emerging single-cell modalities (CITE-seq, multiome, Slide-seq). Comparing traditional statistical methods to embeddings from foundation models trained on millions of cells represents an important emerging direction. Integration of methods across modalities (RNA + protein, RNA + chromatin) would address increasingly common multi-omics experiments.

## 5 Conclusion

This systematic comparison of 16 feature selection methods across three single-cell datasets provides practical guidance for scRNA-seq analysis. We found that method choice significantly impacts selected features, with mean pairwise overlap of only 23.7%, yet pathway-level interpretations remain largely consistent across methods (50–100% pathway overlap). Supervised feature selection methods (ANOVA F-test, Random Forest, Brennecke Method) demonstrated superior recovery of known cell type markers (85–100% for most cell types), while unsupervised approaches (Seurat HVG, CellRanger HVG, variance-based methods) provided broader biological coverage without requiring prior annotations. These supervised methods include both classical statistical tests (ANOVA F-test, Chi-Square test) and machine learning algorithms (Random Forest, Mutual Information), unified by their requirement for labeled data.

Deep learning methods (IntegratedGradients, DeepLIFT, GradientShap) formed a distinct methodological family with high internal consistency (Jaccard 0.6–0.8) and unique gene selections compared to statistical approaches. They showed particularly strong enrichment for immune pathways, suggesting they capture complex biological patterns. However, their longer execution times (1.02–1.18 seconds versus 0.20–0.35 seconds for fast statistical methods) and lower marker recovery rates (71.8–76.1% versus 85–100% for supervised methods) suggest they are best used as complementary approaches rather than primary methods for most applications.

A core set of 1,150 consensus genes selected by at least 50% of methods represents high-confidence features enriched for protein-coding genes (87.6%) rather than technical artifacts. All methods successfully identified relevant biological pathways with high statistical significance, though with varying emphasis on immune response, gene expression, metabolism, and signaling categories.

For practitioners, we recommend CellRanger HVG or Seurat HVG as robust defaults for exploratory analysis, supervised methods when validated cell type markers are the primary goal, and consideration of ensemble approaches combining multiple method families for maximum robustness. The fast execution of most methods (0.20–0.68 seconds for statistical and supervised approaches) means computational efficiency should not drive method selection for typical datasets. Instead, biological goals, availability of annotations, and desired balance between sensitivity and specificity should guide method choice.

The substantial method-specific feature selection observed here, combined with convergent pathway-level interpretations, suggests that biological conclusions in scRNA-seq analysis are more robust than they may initially appear from feature-level comparisons. However, this robustness should not discourage methodological rigor. Transparent reporting of feature selection methods, validation using orthogonal approaches, and consideration of multiple feature sets remain essential for reproducible single-cell genomics.

