## supplementary materials for "Comparative Analysis of Feature Selection Methods for Single-Cell RNA Sequencing Data"

Figure S1

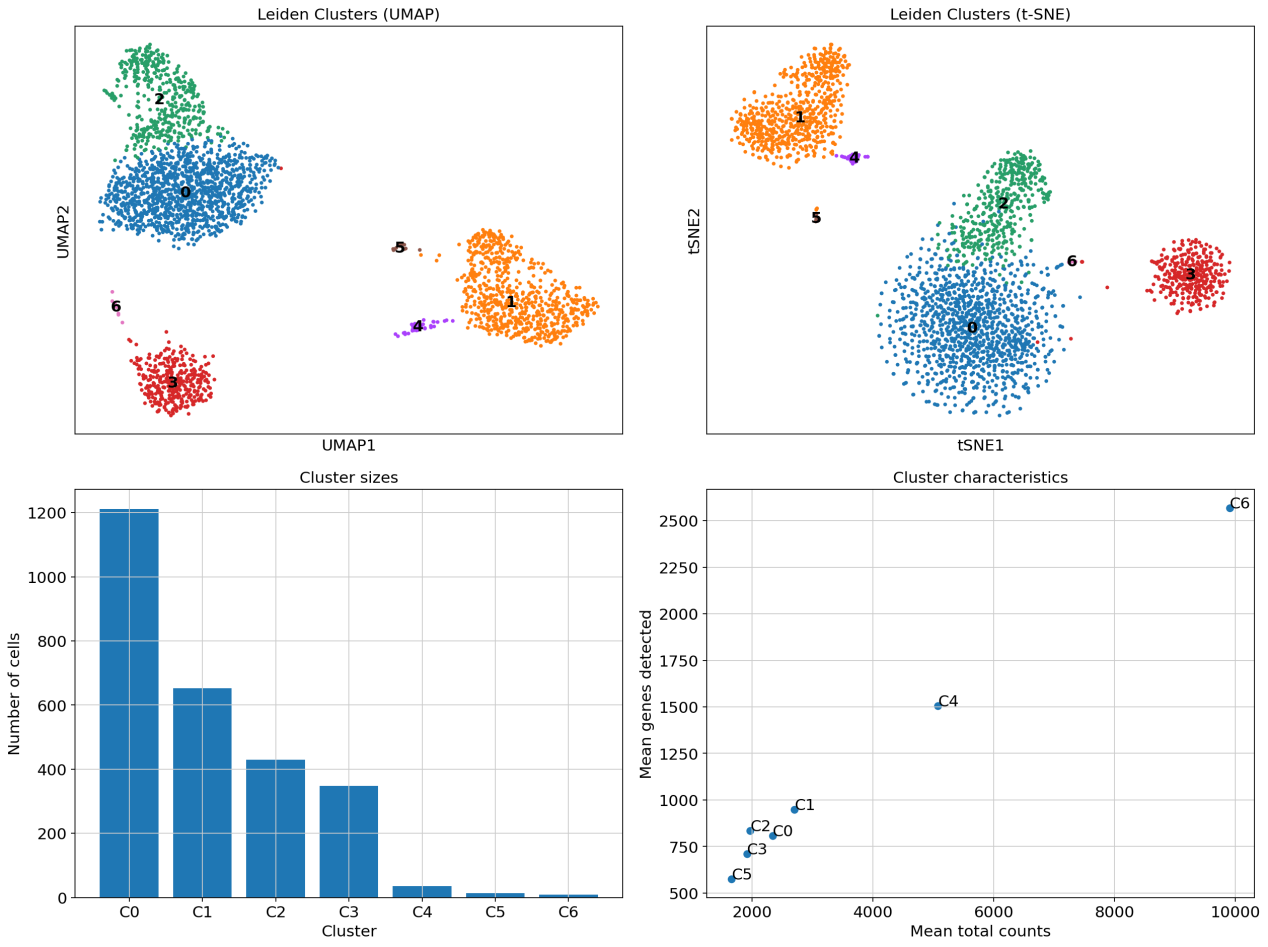

**Figure S1. Quality control metrics for PBMC 3K dataset.** Distribution of key quality control parameters across 2,700 peripheral blood mononuclear cells. (Top row) Histograms showing distribution of total UMI counts per cell, number of genes detected per cell, and mitochondrial gene percentage. Red dashed lines indicate median values. (Bottom row) Scatter plots examining relationships between total UMI counts, number of genes detected, and mitochondrial gene percentage to identify potential low-quality cells or doublets.

Figure S2

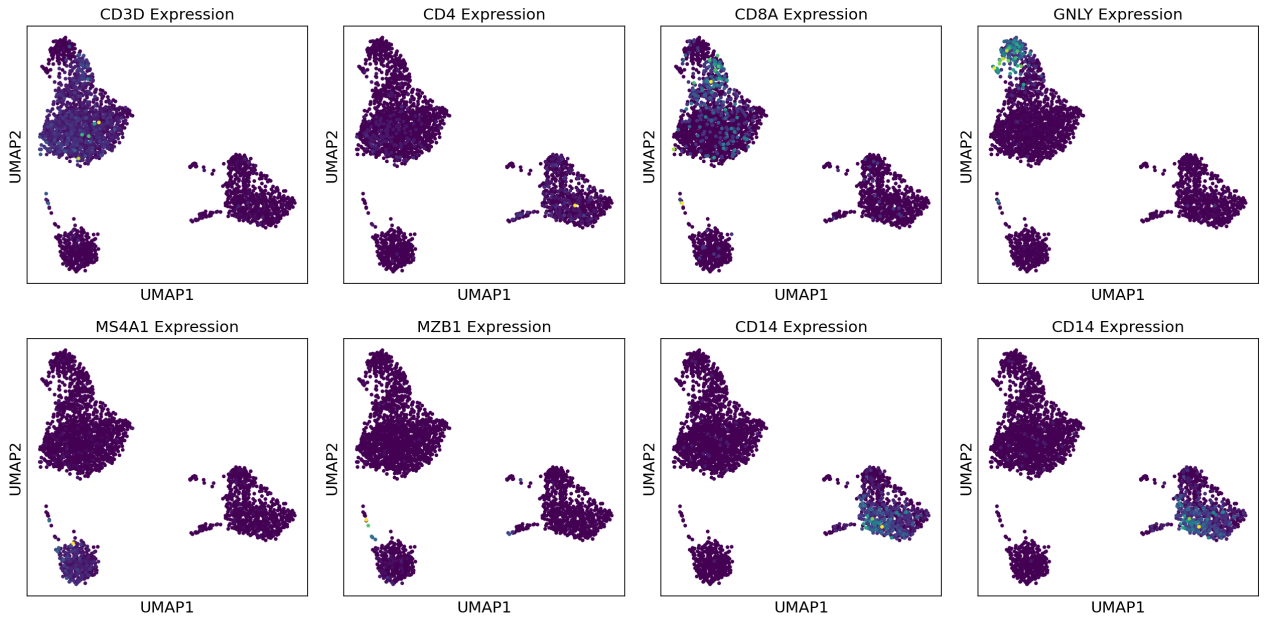

**Figure S2. Clustering analysis of PBMC 3K dataset.** Unsupervised clustering reveals distinct immune cell populations. (Top left) UMAP projection colored by Leiden cluster assignments showing separation of major cell populations. (Top right) t-SNE projection with corresponding cluster labels. (Bottom left) Bar plot showing the number of cells in each identified cluster. (Bottom right) Scatter plot of cluster characteristics showing mean total counts versus mean genes detected per cluster, with cluster labels annotated.

Figure S3

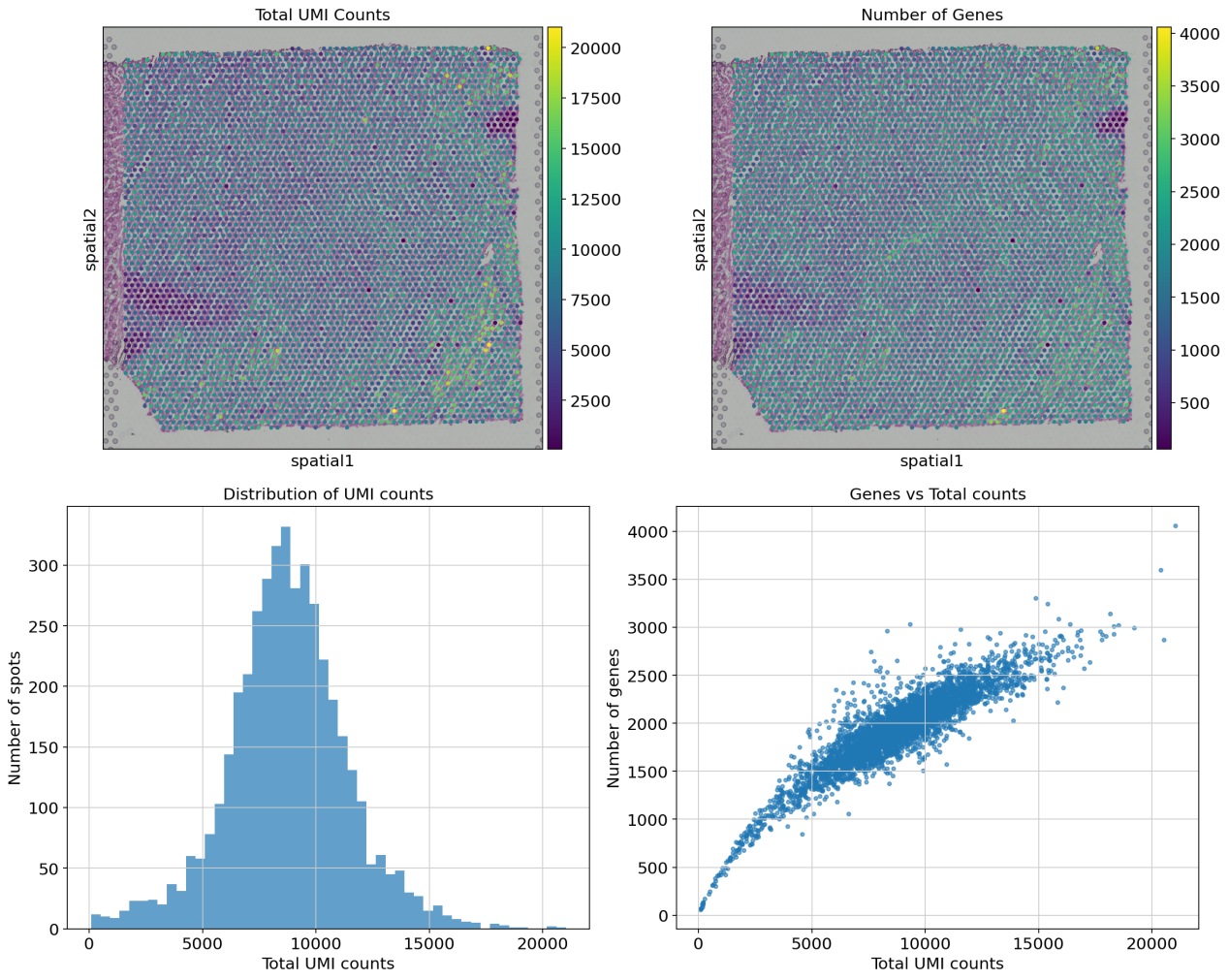

**Figure S3. Spatial quality metrics for Visium human heart dataset.** Quality control analysis of spatial transcriptomics data from human heart tissue. (Top row) Spatial plots overlaid on tissue histology showing distribution of total UMI counts per spot and number of genes detected per spot, revealing regional variation in RNA capture efficiency. (Bottom row) Distribution histogram of UMI counts across all spots and scatter plot examining the relationship between total counts and gene detection.

Figure S4

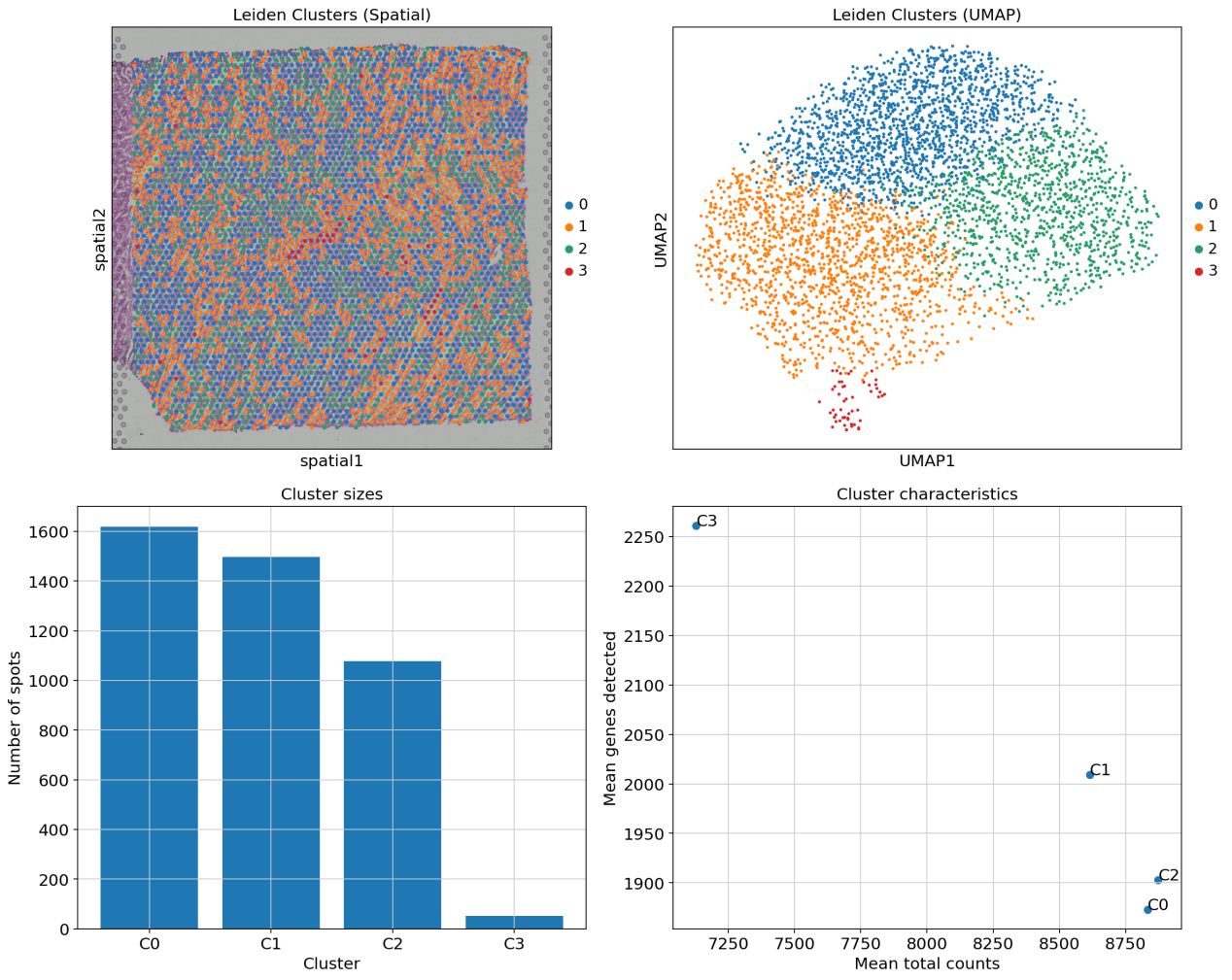

Figure S5

**Figure S4. Spatial clustering analysis of Visium human heart dataset.** Unsupervised clustering identifies distinct cardiac tissue regions. (Top left) Leiden clusters visualized on tissue spatial coordinates overlaid on histology image, revealing anatomically coherent regions. (Top right) UMAP projection of spots colored by cluster assignment. (Bottom left) Bar chart showing the number of spots per cluster. (Bottom right) Scatter plot of mean transcriptomic characteristics (total counts vs. genes detected) for each cluster.

Figure S5

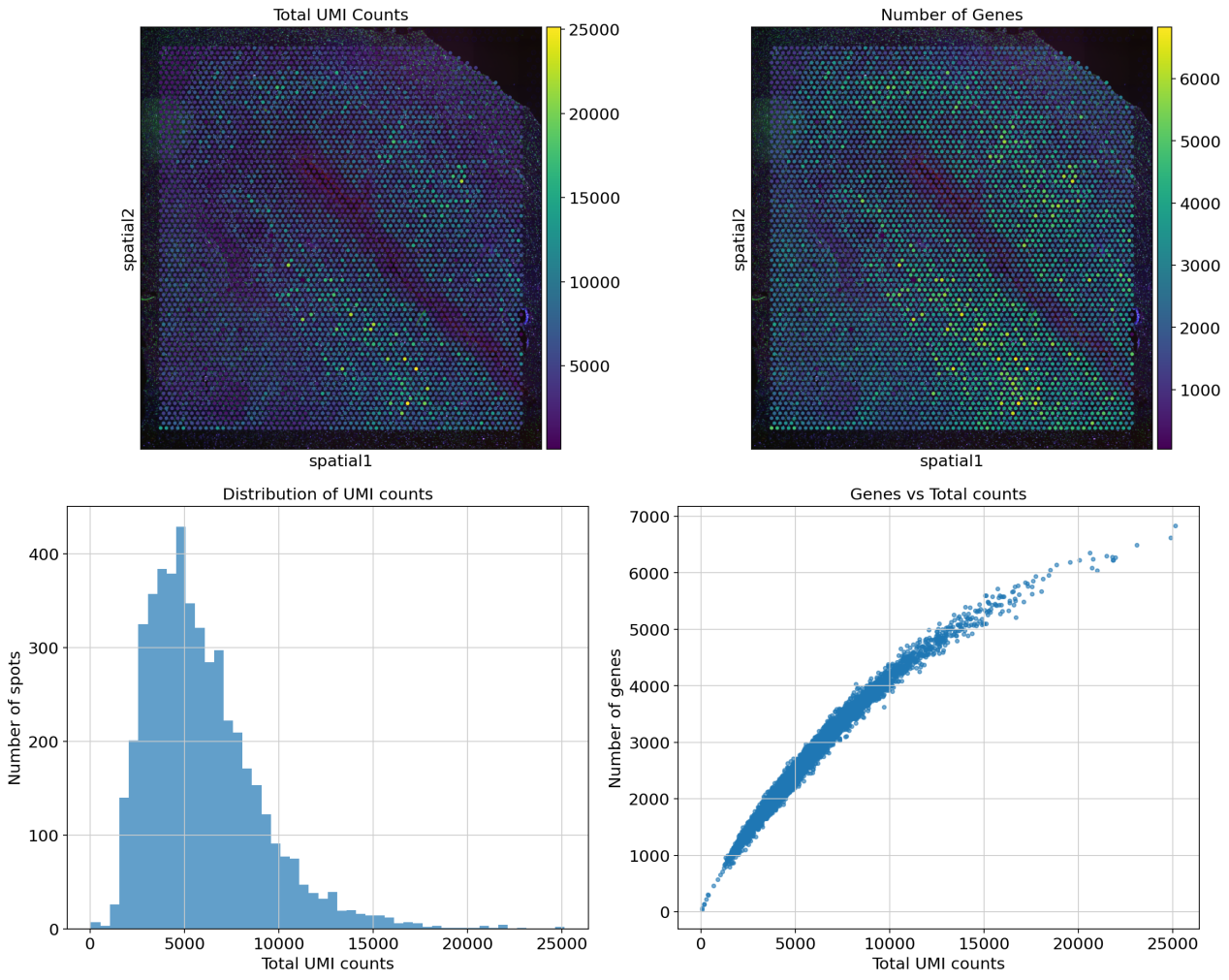

**Figure S5. Spatial quality metrics for Visium human brain dataset.** Quality control analysis of spatial transcriptomics data from human brain tissue section. (Top row) Spatial visualization of total UMI counts and genes detected per spot overlaid on brain tissue histology, showing regional differences in transcriptional activity. (Bottom row) Distribution histogram of total UMI counts and scatter plot of genes detected versus total counts across all spatial spots.

Figure S6

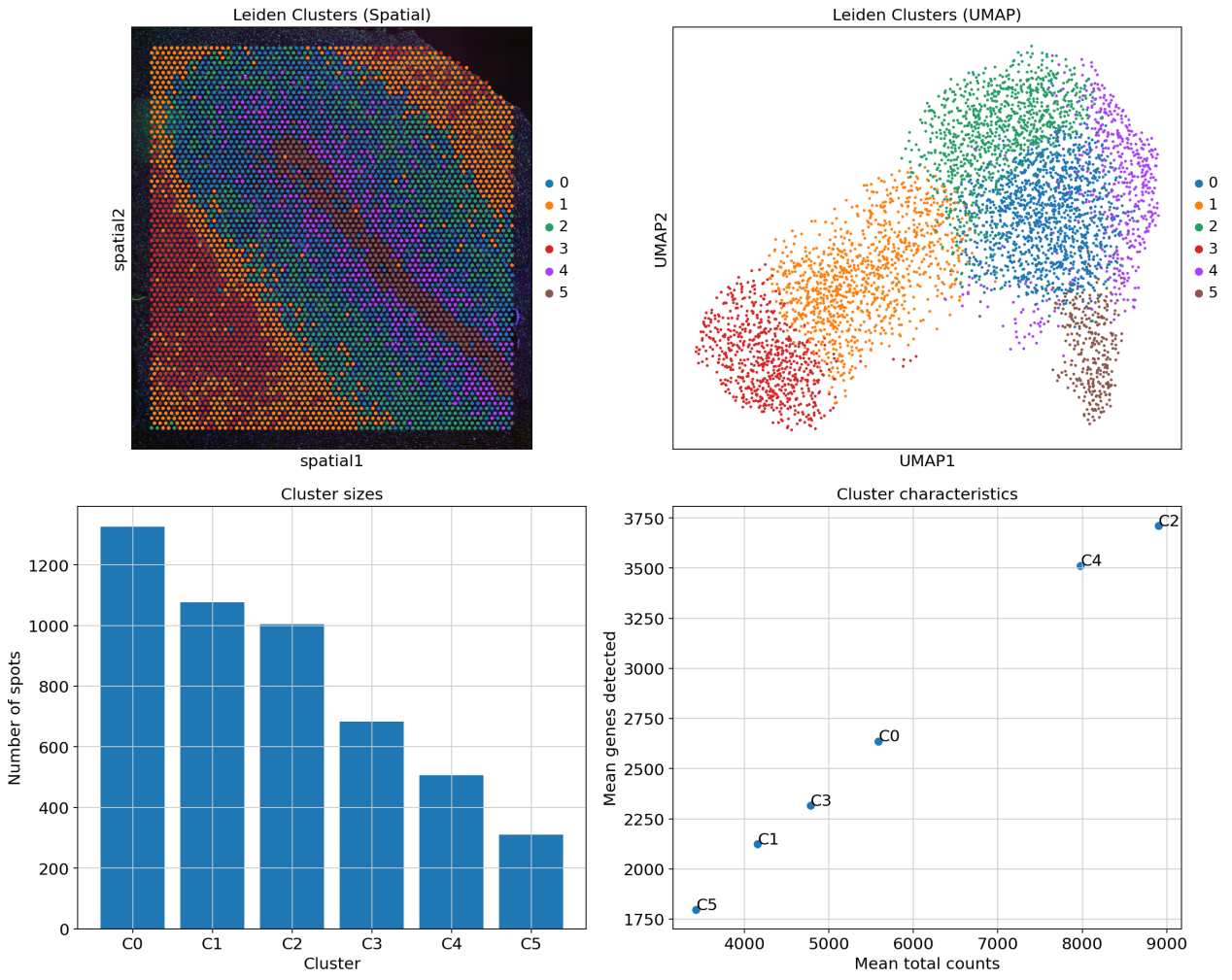

**Figure S6. Spatial clustering analysis of Visium human brain dataset.** Unsupervised clustering reveals distinct brain anatomical regions. (Top left) Leiden clusters mapped onto tissue spatial coordinates with histology background, demonstrating spatially contiguous regions corresponding to brain anatomy. (Top right) UMAP embedding of spots colored by cluster identity. (Bottom left) Distribution of spots across identified clusters. (Bottom right) Mean transcriptomic profiles of each cluster showing relationship between UMI counts and gene detection.

**Table S1. Marker Recovery**

| **Method** | **Cell Type** | **Total Markers** | **Recovered** | **Recovery %** | **Recovered Genes** |
| --- | --- | --- | --- | --- | --- |
| Seurat_HVG | B_cells | 4 | 4 | 100 | CD19, MS4A1, CD79A, CD79B |
| CellRanger_HVG | B_cells | 4 | 4 | 100 | CD19, MS4A1, CD79A, CD79B |
| DeepLIFT | B_cells | 4 | 4 | 100 | CD19, MS4A1, CD79A, CD79B |
| GradientShap | B_cells | 4 | 4 | 100 | CD19, MS4A1, CD79A, CD79B |
| Seurat_v3_HVG | B_cells | 4 | 3 | 75 | MS4A1, CD79A, CD79B |
| Pearson_Residuals | B_cells | 4 | 3 | 75 | MS4A1, CD79A, CD79B |
| Brennecke_Method | B_cells | 4 | 3 | 75 | MS4A1, CD79A, CD79B |
| Variance_Based | B_cells | 4 | 3 | 75 | MS4A1, CD79A, CD79B |
| Gini_Coefficient | B_cells | 4 | 3 | 75 | MS4A1, CD79A, CD79B |
| PCA_Loadings | B_cells | 4 | 3 | 75 | MS4A1, CD79A, CD79B |
| IntegratedGradients | B_cells | 4 | 3 | 75 | MS4A1, CD79A, CD79B |
| Chi2_Selection | B_cells | 4 | 1 | 25 | CD79A |
| ANOVA_F_Test | B_cells | 4 | 1 | 25 | CD79A |
| Random_Forest | B_cells | 4 | 1 | 25 | CD79A |
| Mutual_Information | B_cells | 4 | 0 | 0 | None |
| CV_Based | B_cells | 4 | 0 | 0 | None |
| Seurat_HVG | Dendritic | 3 | 3 | 100 | FCER1A, CST3, CLEC10A |
| Seurat_v3_HVG | Dendritic | 3 | 3 | 100 | FCER1A, CST3, CLEC10A |
| CellRanger_HVG | Dendritic | 3 | 3 | 100 | FCER1A, CST3, CLEC10A |
| Pearson_Residuals | Dendritic | 3 | 3 | 100 | FCER1A, CST3, CLEC10A |
| Gini_Coefficient | Dendritic | 3 | 3 | 100 | FCER1A, CST3, CLEC10A |
| PCA_Loadings | Dendritic | 3 | 3 | 100 | FCER1A, CST3, CLEC10A |
| Brennecke_Method | Dendritic | 3 | 2 | 66.7 | FCER1A, CST3 |
| Chi2_Selection | Dendritic | 3 | 1 | 33.3 | CST3 |
| Variance_Based | Dendritic | 3 | 1 | 33.3 | CST3 |
| ANOVA_F_Test | Dendritic | 3 | 1 | 33.3 | CST3 |
| Random_Forest | Dendritic | 3 | 1 | 33.3 | CST3 |
| IntegratedGradients | Dendritic | 3 | 1 | 33.3 | CST3 |
| DeepLIFT | Dendritic | 3 | 1 | 33.3 | CST3 |
| GradientShap | Dendritic | 3 | 1 | 33.3 | CST3 |
| Mutual_Information | Dendritic | 3 | 0 | 0 | None |
| CV_Based | Dendritic | 3 | 0 | 0 | None |
| CellRanger_HVG | Monocytes | 5 | 5 | 100 | CD14, LYZ, S100A8, S100A9, FCGR3A |
| Pearson_Residuals | Monocytes | 5 | 5 | 100 | CD14, LYZ, S100A8, S100A9, FCGR3A |
| Brennecke_Method | Monocytes | 5 | 5 | 100 | CD14, LYZ, S100A8, S100A9, FCGR3A |
| Variance_Based | Monocytes | 5 | 5 | 100 | CD14, LYZ, S100A8, S100A9, FCGR3A |
| Gini_Coefficient | Monocytes | 5 | 5 | 100 | CD14, LYZ, S100A8, S100A9, FCGR3A |
| PCA_Loadings | Monocytes | 5 | 5 | 100 | CD14, LYZ, S100A8, S100A9, FCGR3A |
| IntegratedGradients | Monocytes | 5 | 5 | 100 | CD14, LYZ, S100A8, S100A9, FCGR3A |
| DeepLIFT | Monocytes | 5 | 5 | 100 | CD14, LYZ, S100A8, S100A9, FCGR3A |
| GradientShap | Monocytes | 5 | 5 | 100 | CD14, LYZ, S100A8, S100A9, FCGR3A |
| Seurat_HVG | Monocytes | 5 | 4 | 80 | LYZ, S100A8, S100A9, FCGR3A |
| Seurat_v3_HVG | Monocytes | 5 | 4 | 80 | LYZ, S100A8, S100A9, FCGR3A |
| Mutual_Information | Monocytes | 5 | 1 | 20 | CD14 |
| Chi2_Selection | Monocytes | 5 | 0 | 0 | None |
| CV_Based | Monocytes | 5 | 0 | 0 | None |
| ANOVA_F_Test | Monocytes | 5 | 0 | 0 | None |
| Random_Forest | Monocytes | 5 | 0 | 0 | None |
| CellRanger_HVG | NK_cells | 4 | 4 | 100 | NKG7, GNLY, KLRD1, NCAM1 |
| Gini_Coefficient | NK_cells | 4 | 3 | 75 | NKG7, GNLY, KLRD1 |
| PCA_Loadings | NK_cells | 4 | 3 | 75 | NKG7, GNLY, KLRD1 |
| IntegratedGradients | NK_cells | 4 | 3 | 75 | NKG7, GNLY, KLRD1 |
| DeepLIFT | NK_cells | 4 | 3 | 75 | NKG7, GNLY, KLRD1 |
| GradientShap | NK_cells | 4 | 3 | 75 | NKG7, GNLY, KLRD1 |
| Seurat_HVG | NK_cells | 4 | 2 | 50 | NKG7, GNLY |
| Seurat_v3_HVG | NK_cells | 4 | 2 | 50 | NKG7, GNLY |
| Pearson_Residuals | NK_cells | 4 | 2 | 50 | NKG7, GNLY |
| Brennecke_Method | NK_cells | 4 | 2 | 50 | NKG7, GNLY |
| Variance_Based | NK_cells | 4 | 2 | 50 | NKG7, GNLY |
| Chi2_Selection | NK_cells | 4 | 1 | 25 | NKG7 |
| ANOVA_F_Test | NK_cells | 4 | 1 | 25 | NKG7 |
| Random_Forest | NK_cells | 4 | 1 | 25 | NKG7 |
| Mutual_Information | NK_cells | 4 | 0 | 0 | None |
| CV_Based | NK_cells | 4 | 0 | 0 | None |
| Variance_Based | T_cells | 7 | 7 | 100 | CD3D, CD3E, CD3G, IL7R, CD4, CD8A, CD8B |
| PCA_Loadings | T_cells | 7 | 7 | 100 | CD3D, CD3E, CD3G, IL7R, CD4, CD8A, CD8B |
| IntegratedGradients | T_cells | 7 | 7 | 100 | CD3D, CD3E, CD3G, IL7R, CD4, CD8A, CD8B |
| DeepLIFT | T_cells | 7 | 7 | 100 | CD3D, CD3E, CD3G, IL7R, CD4, CD8A, CD8B |
| GradientShap | T_cells | 7 | 7 | 100 | CD3D, CD3E, CD3G, IL7R, CD4, CD8A, CD8B |
| Brennecke_Method | T_cells | 7 | 6 | 85.7 | CD3D, CD3E, IL7R, CD4, CD8A, CD8B |
| Gini_Coefficient | T_cells | 7 | 6 | 85.7 | CD3D, CD3E, CD3G, IL7R, CD8A, CD8B |
| Pearson_Residuals | T_cells | 7 | 4 | 57.1 | CD3D, CD3E, IL7R, CD8B |
| CellRanger_HVG | T_cells | 7 | 2 | 28.6 | CD8A, CD8B |
| Mutual_Information | T_cells | 7 | 1 | 14.3 | IL7R |
| Seurat_HVG | T_cells | 7 | 0 | 0 | None |
| Seurat_v3_HVG | T_cells | 7 | 0 | 0 | None |
| Chi2_Selection | T_cells | 7 | 0 | 0 | None |
| CV_Based | T_cells | 7 | 0 | 0 | None |
| ANOVA_F_Test | T_cells | 7 | 0 | 0 | None |
| Random_Forest | T_cells | 7 | 0 | 0 | None |
